# Alzheimer’s disease causes bone marrow myelopoiesis dysfunction

**DOI:** 10.1101/2025.10.06.680030

**Authors:** Miguel Angel Abellanas, Leyre Basurco, Maitreyee Purnapatre, Chiara Burgaletto, Giulia Castellani, Sarah Phoebeluc Colaiuta, Javier Maria Peralta-Ramos, Angham Ibraheem, Sama Murad, Paola Antonello, Mariangeles Kovacs, Yuliya Androsova, Bar Nathansohn, Hannah Partney, Liora Cahalon, Rafael Valdes-Mas, Joseph M. Josephides, Tomer M. Salame, Maria Espelosin, Mar Cuadrado-Tejedor, Ana Garcia-Osta, Aleksandra Deczkowska, Michal Schwartz

## Abstract

Accumulating evidence suggests that both innate and adaptive immunity play crucial roles in combating Alzheimer’s disease (AD). Specifically, enhancing the homing of monocyte-derived macrophages to the affected brain has been shown to reduce local inflammation, decrease proteinopathy, rescue neurons, and mitigate cognitive decline. However, the factors limiting their spontaneous recruitment remain unclear. Using multi-omics techniques, we identified impaired myelopoiesis and monocyte development in both mice and AD patients. While not the primary cause of the disease, this impairment is associated with disease progression. In the 5xFAD mouse model, monocyte differentiation was found to be disrupted due to a maladaptive bone marrow (BM) response, driven by type I interferon (IFN-I) signaling. A similar phenotype was found in circulating monocytes from AD patients compared to healthy controls. Blocking IFN-I with monoclonal antibodies or using chimeric AD mice with BM from mice lacking the IFN-I receptor (IFNAR1) alleviated myelopoiesis dysfunction, normalized monocyte phenotypes, and reduced cognitive impairment. These improvements in myeloid function were accompanied by an increased homing of monocyte-derived macrophages in the AD brain. Our results reveal an unexpected dysfunction in BM myelopoiesis in the context of neurodegeneration and support the emerging concept that neurodegenerative diseases are not solely brain-centric.

## Introduction

Alzheimer’s disease (AD) was considered for decades to be a brain-centric disease. Currently, there is a growing appreciation for the role of the immune component in disease progression and, therefore, also for its therapeutic potential target. In this context, resident and infiltrating myeloid cells are crucial players in the immune response to AD^1^. In mouse models of AD, for example, early stages of the disease are associated with TREM2-dependent activated microglia, the so-called disease-associated microglia (DAM)^2,3^. This phenotype is suggested to contain the amyloid pathology^4^. In contrast, however, in the course of disease deterioration, microglia seem to be detrimental^5,6^ and eventually develop a senescent phenotype that further worsens cognitive dysfunction^7^. Therefore, the limited ability of microglial cells to cope with AD pathology challenges the understanding of these cells as the sole defenders of brain health^1^.

In contrast to the microglia, infiltrating monocyte-derived macrophages were shown to be beneficial under acute and chronic brain pathologies^8–13^, including AD pathology^14–18^. The genetic blockade of their recruitment to the brain through C-C chemokine receptor 2 (CCR2) accelerates disease manifestation^14^, while non-classical monocytes were shown to remove amyloid from the blood vessels, limiting vascular amyloidosis^18^. Furthermore, the adoptive transfer of young and healthy monocytes is protective in a familial mouse model of AD^16^, suggesting that proper monocyte function is necessary to limit AD pathology. Therefore, given the need for monocyte-derived macrophages, it is puzzling why their recruitment does not occur spontaneously in AD and what mechanisms prevent this. Considering the dynamic nature of these cells, one possible explanation is that their differentiation in the bone marrow (BM) is affected by AD, as happens in acute neurological injuries and some neurodegenerative diseases^19–22^. The phenotype of these cells and their protective functions could be impaired in AD, as a recent multiomic atlas of peripheral blood mononuclear cells (PBMC) suggests^23^. It was found that among all cell types, monocytes are the ones mostly affected by the disease, showing a transcriptomic and epigenetic bias towards a proinflammatory state^23^.

Here, we discovered that the factor limiting the recruitment of resolving monocytes in AD stems from the dysfunction of BM myelopoiesis. We demonstrate that hematopoietic stem cells (HSC) are reduced in the BM throughout disease progression in the 5xFAD mouse model of familial AD. This loss of HSC significantly impacts monopoiesis and is driven by circulating type I interferon (IFN-I) signaling. As a corollary, chronic exposure of 5xFAD BM to IFN-I further impaired monocyte maturation, which resulted in a reduction of both non-classical patrolling monocytes and infiltrating macrophages in the brain. The phenotype of the classical monocytes in the brain overlapped with those in the blood and the BM, suggesting a conserved phenotype between these compartments. This pathological phenotype in classical monocytes was also found in another animal model of AD and PBMCs from AD patients. Moreover, blocking IFN-I signaling in the periphery preserved HSC quiescence, enhanced BM myelopoiesis, and delayed disease progression, prevented memory loss, and boosted macrophage differentiation within the brain. These findings suggest that circulating IFN-I impairs hematopoiesis in the peripheral BM and accelerates disease progression by preventing monocytes/macrophages from acquiring a resolving phenotype.

## Results

### Alzheimer’s disease is associated with impaired myelopoiesis in the bone marrow

Hematopoiesis is a multi-step process tightly governed by internal and external factors. Long-term (LT) HSC are the most immature stem cells in the BM with the highest differentiation capacity^24,25^, and their maintenance is critical for ensuring proper immune responses^26,27^. To investigate the impact of AD progression on hematopoiesis, we first characterized the different subsets of hematopoietic stem and progenitor cells (HSPC) by flow cytometry in the femur BM, at different ages in 5xFAD mice, compared to age-matched wild-type (WT) littermates (Fig. 1a-c). We defined the HSPC as lineage (Lin)^-^Sca1^+^cKit^+^ (LSK) cells, and the different subtypes of HSPC were defined based on the differential expression of CD48 and CD150 (Fig. 1b, c) as previously described^28^. Among the HSPC subtypes, LT-HSC (Lin^-^Sca1^+^cKit^+^CD150^+^CD48^-^) accumulated in the WT BM as a part of the aging process^29^ (Fig. 1d, e). Yet, at 12-14-month-old 5xFAD, LT-HSC accumulation was significantly reduced in the BM relative to age-matched WT (Fig. 1d, e). Importantly, this reduction was specific to LT-HSC, and not due to a general loss of LSK (Extended Data Fig. 1a).

**Fig. 1:**
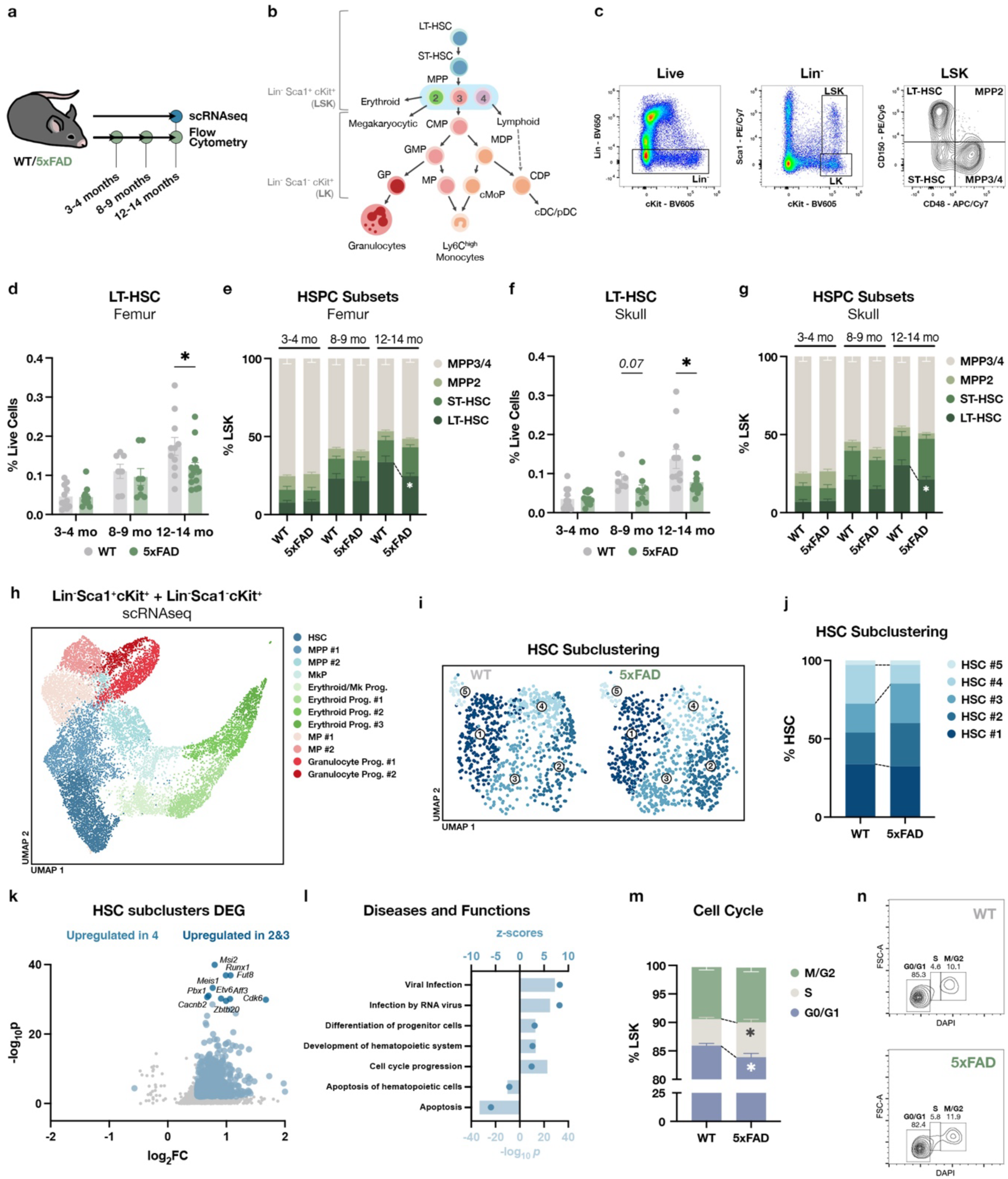
Hematopoietic stem cell quiescence is lost in the 5xFAD mouse model of familial AD. **a**, WT and 5xFAD mice were euthanized at different time points to study BM cell composition by flow cytometry and single-cell RNAseq (scRNAseq) to study HSPC from 12-14-month-old animals. **b**, Scheme representing myelopoiesis^28,33–36^. **c**, Gating strategy to analyze HSPC subpopulations in the BM by flow cytometry. **d**, LT-HSC abundance in the femur BM of WT and 5xFAD animals at different time points analyzed by flow cytometry. Mann-Whitney test. **e**, Proportions of different HSPC subsets in the femur BM of WT and 5xFAD mice at different time points analyzed by flow cytometry. Unpaired t-test. **f**, LT-HSC abundance in the skull BM of WT and 5xFAD animals at different time points analyzed by flow cytometry. Mann-Whitney test. **g**, Proportions of different HSPC subsets in the skull BM of WT and 5xFAD mice at different time points analyzed by flow cytometry. Unpaired t-test. **h**, UMAP showing LSK and LK cells analyzed by scRNAseq from the femur BM of 12-14-month-old WT and 5xFAD mice (WT, *N* = 3; 5xFAD, *N* = 4; all were females). **i**, UMAP showing the subclusterization of the HSC cluster. **j**, Proportions of the different HSC subclusters in the BM of WT and 5xFAD mice. **k**, Differentially expressed genes (DEG) between more abundant HSC clusters in 5xFAD mice (clusters 2 and 3) versus the HSC cluster more abundant in WT mice (cluster 4). Genes highlighted in pale blue have log_2_FC > 0.5 or < −0.5 and p-value < 0.01, and those in dark blue are the top 10 genes. **l**, Ingenuity Pathway Analysis (IPA) ‘Diseases and Functions’ enriched in the DEG from the different HSC subclusters, selected among the ones with z-score > 2 or < −2, and p-value < 0.01. **m**, Flow cytometry analysis of the cell cycle state of LSK from the BM of 12-month-old WT and 5xFAD animals. Data are represented as mean ± SEM. Unpaired t-test. **n**, Contour plot showing DAPI staining of WT and 5xFAD LSK.

The skull BM is highly influenced by the brain signals, since microchannels were shown to directly connect the skull BM to the CNS borders^30,31^. Furthermore, because the skull BM vascular niche was shown to be largely protected against major hallmarks of ageing^32^, we questioned whether the observed changes with aging in LT-HSC in the femur of WT also occur in skull BM. We separately analyzed skull BM from 5xFAD mice and found a similar phenotype to the femur BM; again, while LT-HSC accumulation in the skull BM increased over time in WT mice, this age-associated increase was not observed in 5xFAD skull BM (Fig. 1f, g). Particularly, there was a trend in the reduction of LT-HSC in the 5xFAD skull BM at earlier timepoints (8-9 months), which became statistically significant at the same timepoint as the femur BM (12-14 months) (Fig. 1f, g). This suggests that in 5xFAD mice, the signals causing the reduction in LT-HSC affected both the skull and femur BM similarly.

Since the LT-HSC have the potential to differentiate into all the immune cell subsets and their progenitors, including the myeloid progenitors, we further explored this reduction and the phenotype of the LT-HSC. In particular, HSPC can differentiate into various myeloid progenitors, with varying degrees of commitment to granulocytic, monocytic, or dendritic cell lineages (Fig. 1b)^28,33–36^. This broad group of myeloid-committed progenitors can be detected by flow cytometry analysis as Lin^-^Sca1^-^cKit^+^ (LK) (Fig. 1 b, c). We sorted LSK and LK cells from the femur BM of 12-month-old WT and 5xFAD mice and performed single-cell RNA sequencing (scRNAseq) analysis employing the 10X Genomics platform. Using initial unsupervised clustering and Uniform Manifold Approximation and Projection (UMAP), we identified clusters of cells expressing gene markers typical of HSC, multipotent progenitors (MPP), and erythroid and myeloid progenitors (Fig. 1h, Extended Data Fig. 1c). We then subclustered the HSC and obtained five different groups (Fig. 1i). While cells present in subclusters 1 and 5 expressed genes characteristic of short-term HSC (ST-HSC) and MPP such as *Cd34* and *Flt3* (Extended Data Fig. 1d), cells in subclusters 2-4 expressed gene markers including *Procr* and *Fgd5*, characteristic of LT-HSC (Extended Data Fig. 1d). Interestingly, we observed a reorganization of the LT-HSC subclusters in the 5xFAD mice compared to the WT (Fig. 1i, j). While the majority of LT-HSC in the WT BM were found in cluster number 4, in the 5xFAD BM, there was a reduction of this cluster in favor of clusters 2 and 3 (Fig. 1j). To further understand this shift in population dynamics, we compared the differentially expressed genes (DEG) between clusters 2-3 and cluster 4. We identified a large number of DEG upregulated in clusters 2 and 3, in the 5xFAD versus WT mice (Fig. 1k). Using Ingenuity Pathway Analysis (IPA), we identified functions that were upregulated and associated with this list of genes, such as “Viral infection”, “Differentiation of progenitor cells” and “Cell cycle progression” (Fig. 1l). We validated the loss of quiescent cells in HSPCs within the 5xFAD bone marrow by analyzing their cell cycle state using flow cytometry. Significant cell cycle progression was observed in the 5xFAD HSPC, characterized by an increase in cells in the S phase and a reduction in the G0/G1 phase (Fig. 1m, n). While we could not detect many DEG comparing the ST-HSC from 5xFAD BM to WT BM, we detected a large number of DEG when analyzing the MPP clusters. These MPPs exhibited a gene expression profile associated with viral response pathways (Extended Data Fig. 1e, f). They also showed reduced expression of *Irf8*, a transcription factor critical for myeloid differentiation (Extended Data Fig. 1e, f), suggesting impaired myeloid potential. Taken together, these findings indicate that LT-HSC in 5xFAD mice exhibit a viral-like response, accompanied by their reduced abundance and quiescence.

### Impaired myelopoiesis is associated with myeloid cell dysfunction in an IFN-I-dependent mechanism

Chronic viral infections and interferons (IFN) have been shown to induce loss of quiescence and depletion of the LT-HSC compartment^26,37–39^. Moreover, some chronic infections can trigger IFN-I-dependent maladaptive responses in the BM by depleting HSC and monocyte progenitors^26,40^. Next, we examined whether this aberrant phenotype observed in the HSC of 5xFAD mice affects their function and ability to differentiate into monocytes. Using fluorescence-activated cell sorting (FACS), we sorted HSPC from both WT or 5xFAD femur BM at different ages, and cultured them in the presence of monocyte differentiation factors (Fig. 2a). While at the early stage of the disease (e.g., 8-month-old) LSK from 5xFAD showed an increased capacity of monocyte differentiation, a sharp decrease was noticed at 12-14 month-old 5xFAD animals in both femur (Fig. 2b) and skull (Extended Data Fig. 2a). To further verify the age-related impairment of the 5xFAD BM we created chimeric mice, in which we transplanted LSK from either WT or 5xFAD together with GFP^+^ WT competitor cells into irradiated GFP^+^ WT recipients. These chimeric mice allowed us to assess the differentiation capacity of HSPC *in vivo* over a period of 4 months (Fig. 2c). While the degree of chimerism in the BM at the end of the experiment was the same in the recipient mice receiving either WT or 5xFAD LSK (Extended Data Fig. 2b), the production of myeloid cells by the transplanted 5xFAD HSPC was significantly reduced compared to that by the transplanted WT HSPC (Fig. 2d). These findings provide functional validation of the impaired ability of the HSPC from the aged AD BM to properly differentiate into myeloid cells.

**Fig. 2:**
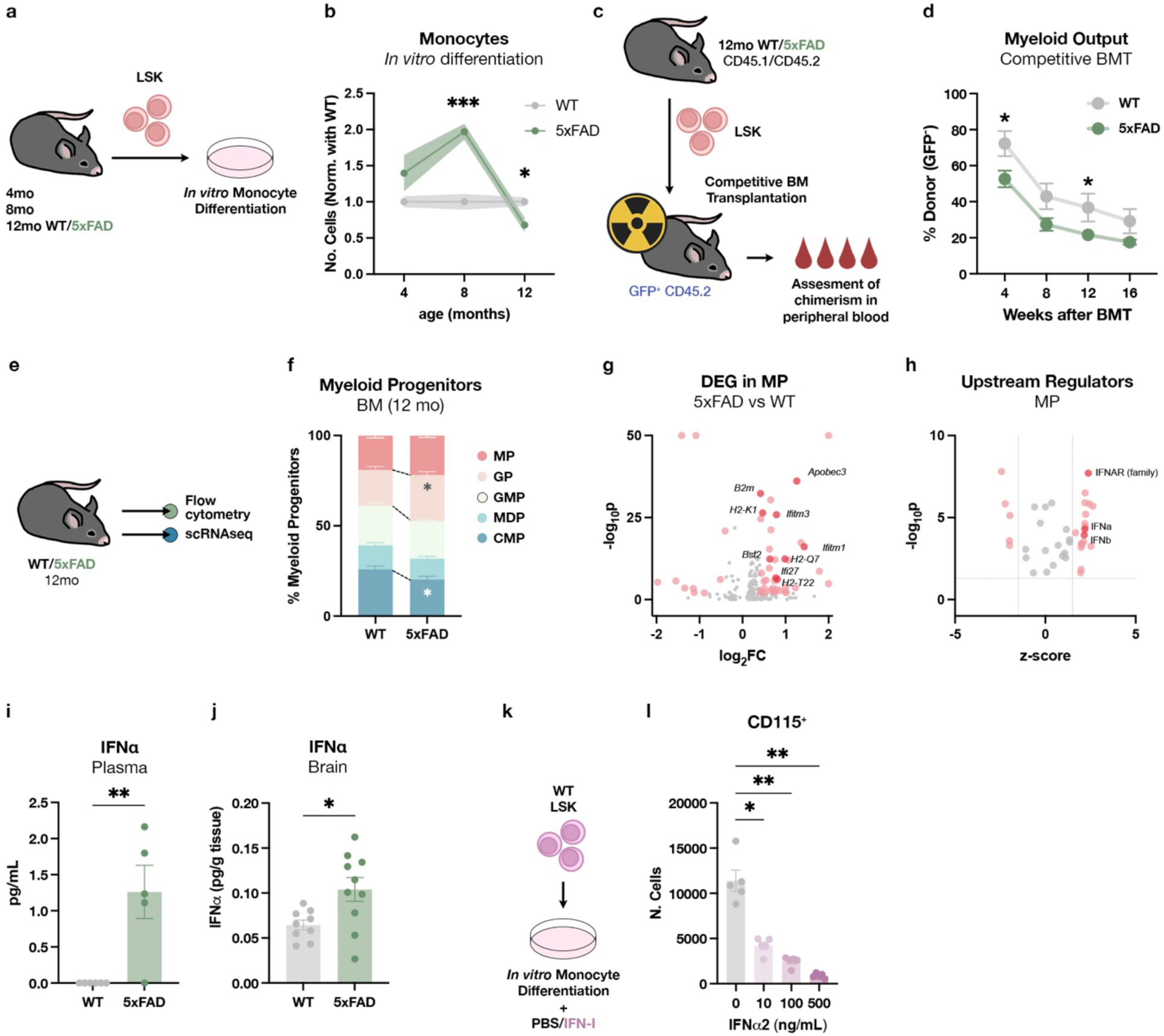
Myelopoiesis is impaired in the 5xFAD BM, showing an IFN-I response. **a**, LSK from 4, 8, and 12-month-old WT and 5xFAD littermates were sorted and cultured in the presence of monocyte-differentiation factors. **b**, Absolute number of differentiated monocytes from WT or 5xFAD LSK after 7 days in culture. Data from 3 independent experiments. Values represent the mean of triplicates normalized with the WT cell cultures. Unpaired t-test at each time point. **c**, LSK from 12-month-old WT or 5xFAD animals were sorted and injected together with total BM GFP^+^ competitor cells into irradiated GFP^+^ recipient mice. Blood was withdrawn from the recipients, every 4 weeks for 16 weeks, and then mice were euthanized. **d**, Flow cytometry analysis of myeloid cell abundance among the GFP^-^ donor cells in the peripheral blood of recipient animals (WT, *N* = 7; 5xFAD, *N* = 9). Unpaired t-test at each time point. **e**, Myeloid progenitors were analyzed in the BM of 12-month-old WT and 5xFAD mice by flow cytometry and scRNAseq. **f**, Myeloid progenitor subsets in the femur BM of WT and 5xFAD mice (WT, *N* = 9; 5xFAD, *N* = 7). Unpaired t-test. **g**, DEG of 5xFAD MP compared to WT MP. Genes highlighted in pink have log_2_FC > 0.4 or < −0.4, and p-value < 0.01, and those in red are related to IFN-I signaling. **h**, Upstream regulators as calculated by IPA. Molecules highlighted in pink have z-score > 1.5 or < −1.5 and p-value < 0.05, and those in red are the molecules related to IFN-I signaling. **i**, IFNα concentration in the plasma collected from 12-month-old WT and 5xFAD animals as analyzed by ELISA (WT, *N* = 6; 5xFAD, *N* = 5). Unpaired t-test. **j**, IFNα concentration in the whole brain collected from 12-month-old WT and 5xFAD animals as analyzed by ELISA (WT, *N* = 9; 5xFAD, *N* = 10). Unpaired t-test. **k**, Sorted LSK were cultured for one week in the presence of IFNα2 and monocyte differentiation factors. **l**, Absolute numbers of monocytes after one week *in vitro* following exposure to different doses of IFNα2. Repeated measures one-way ANOVA followed by Dunnett’s multiple comparisons test.

To further understand the mechanism underlying the impairment of the BM progenitors in the 5xFAD animals, we characterized the myeloid progenitor subsets by flow cytometry and scRNASeq (Fig. 2e). Our analysis revealed a significant loss of the common myeloid progenitors (CMP) and an increase in the granulocyte progenitors (GP) in the BM of 12-month-old 5xFAD compared to age-matched WT mice (Fig. 2f, Extended Data Fig. 2c). This effect was further validated using a knock-in APP mouse model (APP^NL-G-F^), where, even if there were no differences in CMP numbers (Extended Data Fig. 2d), there was a loss of monocyte progenitors (MP) in the femur BM of APP^NL-G-F^ compared to age matched WT (Extended Data Fig. 2e). Taken together, these findings indicate that the myelopoietic capacity of HSPC from the 5xFAD femur BM is impaired at an age at which the disease is fully manifested. By performing differential gene expression analysis of MP from 5xFAD mice compared to their WT counterparts in our single-cell dataset (Fig. 2g), and inferring upstream regulators by IPA, we detected an enrichment of genes modulated by the IFN-I family signaling, including IFN-α and β (Fig. 2h). Taken together, these findings indicate that the myelopoietic capacity of HSPC in two independent animal AD models is impaired. This impairment is pronounced at an age at which the disease is fully manifested, potentially contributing to the limited availability of monocyte-derived macrophages precisely when they are most needed. Of note, the detected impaired phenotype in the BM function seemed to be selective to the myeloid lineage, as we could not detect any significant changes in the proportions of lymphocyte progenitors or major lymphoid populations in the femur BM (Extended Data Fig. 3a-c) or in other peripheral immune tissues at the same age (Extended Data Fig. 3d-f).

We considered that the IFN-I signature detected in our BM transcriptomic analysis might be an outcome of the presence of circulating IFN-I, likely derived from the brain^6^. Using enzyme-linked immunosorbent assay (ELISA), we found increased levels of IFN-α in the plasma of the 5xFAD, which were undetectable in the age-matched WT mice (Fig. 2i). No detectable increase in IFN-α was observed in the bone marrow of 5xFAD mice (Extended Data Fig. 2f). In addition to the plasma, increased levels of IFN-α were observed in the brain of 5xFAD mice (Fig. 2j), pointing to the CNS as the potential primary source of the circulating IFN-α. No increase in IFN-β was detected in any of the compartments tested in 5xFAD mice compared to WT animals (Extended Data Fig. 2g, h).

To gain mechanistic insight into the effect of IFN-I on monopoiesis, we took advantage of our *in vitro* differentiation system (Fig. 2k). By exposing sorted WT LSK to increasing doses of a member of the IFN-α family together with monocyte differentiation factors, we detected a significant dose-dependent reduction in the number of differentiated monocytes after 7 days *in vitro* (Fig. 2l). This effect was reproduced when exposing the cells to another member of the IFN-I family, the IFN-β (Extended Data Fig. 4a-b).

To further validate the effect of IFN-I on monocyte differentiation, we systemically injected polyinosinic:polycytidylic acid (poly I:C), a strong inducer of IFN-I production by mimicking a viral infection, to either WT or IFNAR1-KO mice, which lack the common receptor for IFN-α/β, and were used here as controls (Extended Data Fig. 4c). Under this administration regime, poly I:C was shown to impair LT-HSC quiescence and monocyte progenitor function^37,40^. A significant reduction of non-classical and intermediate monocytes (Ly6C^low^ and Ly6C^int^, respectively) was observed, with a proportional increase in classical monocytes (Ly6C^high^) in the BM following poly I:C treatment (Extended Data Fig. 4d, e). The same poly I:C treatment in transgenic mice lacking IFNAR1 failed to induce these phenotypes (Extended Data Fig. 4e). A similar effect was detected in the blood but not in the spleen (Extended Data Fig. 4f, g), suggesting an impairment initiated in the BM, as the origin of these cells before their release into the bloodstream. Given that non-classical monocytes arise from classical monocytes^41^, we further hypothesized that this rearrangement in monocyte subpopulations could be the outcome of the retention of classical monocytes in the BM.

Monocyte release into the circulation involves a balance between chemoretention and chemoattraction^42^. The C-X-C motif chemokine receptor 4 (CXCR4) restrains the release of immature immune cells in the BM and is a marker of immature monocytes^43^, and it must be downregulated to enable their release into the bloodstream^42^. We therefore assessed levels of CXCR4 expression by BM-derived classical monocytes following poly I:C treatment. Classical monocytes expressed significantly higher levels of CXCR4 in the BM of poly I:C-treated mice (Extended Data Fig. 4h,i), consistent with their increased accumulation in the BM and impaired differentiation to non-classical monocytes (Extended Data Fig. 4e). This increase of CXCR4 expression was also IFN-I-dependent, since the IFNAR-KO animals treated with poly I:C did not show it (Extended Data Fig. 4i).

To further study the phenotype of the classical monocytes following poly I:C treatment, we performed bulk RNAseq and detected a significant shift in the transcriptomic profile in these cells (Extended Data Fig. 4j, k). Analysis by IPA revealed that both up- and downregulated DEGs were associated with increased IFN-α signaling as well as with interferon regulatory factor 3 (IRF3), IRF7, and IRF9; all of them known to mediate IFN-I responses (Extended Data Fig. 4l). Neuroinflammation and IFN-I-related pathways were significantly enriched in classical monocytes from poly I:C-treated mice (Extended Data Fig. 4m). These evidences further confirm the detrimental effect of IFN-I signaling on classical monocyte phenotype, not only by preventing their maturation and release, but by also inducing a bias towards a pro-inflammatory response.

### Monocytes acquire a proinflammatory phenotype in both mouse models and human AD

Since IFN-I impairs monocyte maturation and their release into circulation, we further characterized the mature myeloid compartment in the 5xFAD BM. By flow cytometry, we detected a significant expansion of classical monocytes with the proportional decrease in non-classical monocytes in the femur BM of 12-month-old 5xFAD (Fig. 3a, b), mirroring the IFN-I-dependent phenotype from poly I:C-injected mice (Extended Data Fig. 4e). This accumulation of classical monocytes in the 5xFAD BM was validated in the femur BM of the APP^NL-G-F^ mouse model of AD (Fig. 3c). The significant increase of classical monocytes in the AD BM was accompanied by an increased expression of CXCR4 in these cells (Fig. 3d), similar to the effects induced by the administration of poly I:C through an IFNAR1-dependent mechanism (Extended Data Fig. 4i). The expansion of classical monocytes was also detected in the blood, with the same increase in CXCR4 (Extended Data Fig. 5a, b). On the other hand, this effect was not significant in the spleen (Extended Data Fig. 5c, d), supporting the effects in the BM as a primary event in the monocytic alterations. In the blood of APP^NL-G-F^ mice, we further validated the expansion of classical monocytes with the decrease in non-classical monocytes (Extended Data Fig. 5e). Altogether, these findings support an accumulation of classical monocytes in the BM of AD mouse models, with a poor differentiation and maturation as suggested by the increase in CXCR4 and the reduction of non-classical monocytes.

**Fig. 3:**
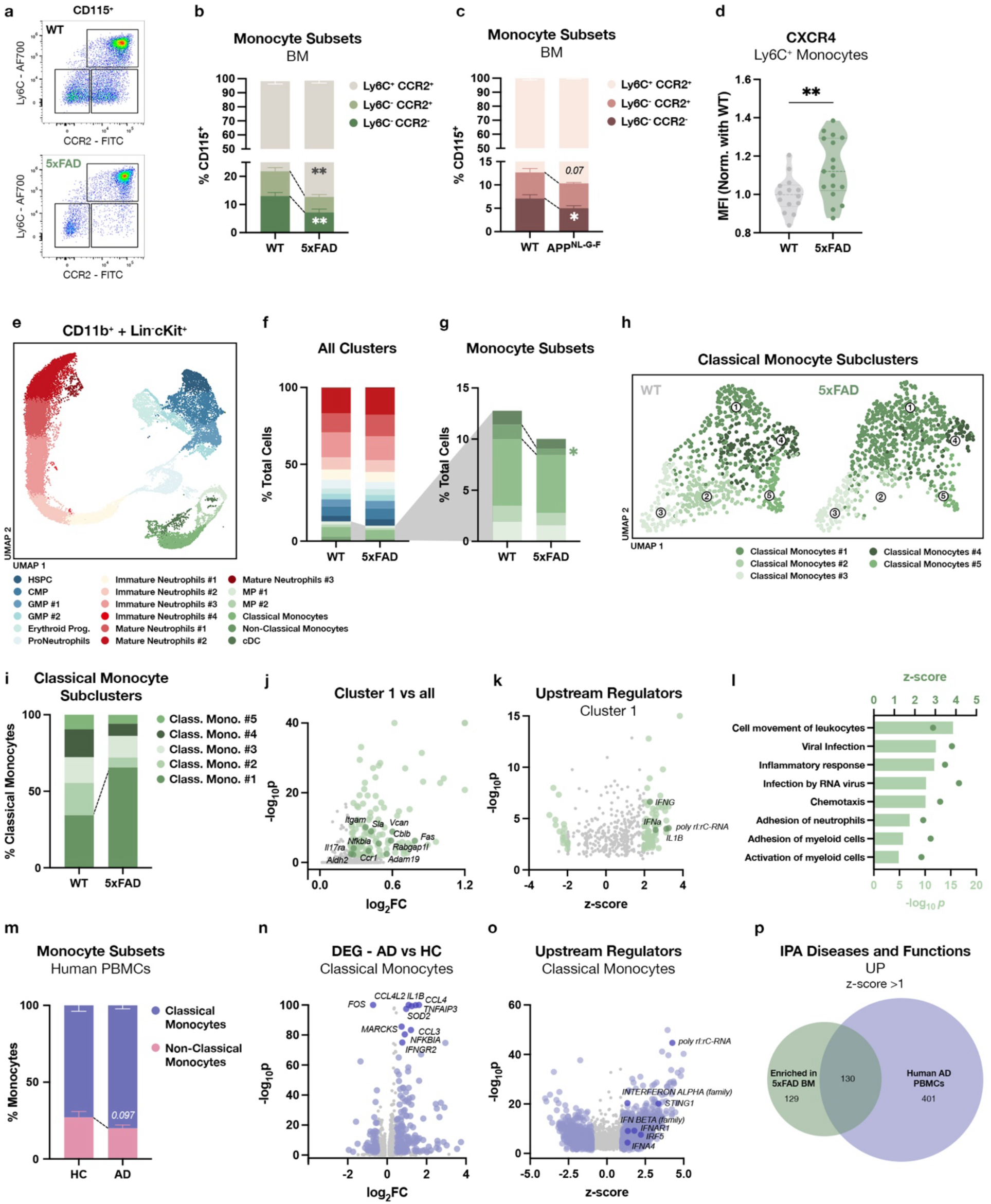
Monocyte phenotype is affected in the periphery of 5xFAD animals and AD patients. **a**, Gating strategy for monocyte subset analysis by flow cytometry. **b**, Monocyte subsets in the BM of 12-month-old WT and 5xFAD mice (WT, *N* = 9; 5xFAD, *N* = 9). Unpaired t-test for each subtype between WT and 5xFAD. **c**, Monocyte subsets in the BM of 20-month-old WT and APP^NL-G-F^ mice (WT, *N* = 5; 5xFAD, *N* = 8). Unpaired t-test for each subtype between WT and APP^NL-G-F^. **d**, Median fluorescence intensity (MFI) of CXCR4 expressed by classical monocytes in the BM of 12-month-old WT and 5xFAD mice (WT, *N* = 14; 5xFAD, *N* = 17). Data were normalized relative to the average MFI of the WT sample in each independent experiment. Unpaired t-test. **e**, UMAP showing clusters generated by scRNAseq analysis of CD11b^+^ and Lineage^-^cKit^+^ sorted from the BM of 12-month-old WT or 5xFAD mice (WT, *N* = 4; 5xFAD, *N* = 4; all were females). **f**, Proportions of the different clusters in the BM of WT and 5xFAD animals. **g**, Proportions of the different monocyte and monocyte progenitor clusters. Unpaired t-test. **h**, UMAP shows the subclusterization of the classical monocyte subclusters. **i**, Proportions of the different classical monocyte subclusters in the BM of WT and 5xFAD mice. **j**, DEG in cluster 1 compared to the other classical monocyte clusters. Genes highlighted in light green have log_2_FC > 0.25 or < −0.25 and p-value < 0.01, and those in dark green are related to IFN-I signaling. **k**, Upstream regulators as calculated by IPA. Molecules highlighted in light green have a z-score > 2 or < −2 and a p-value < 0.05. **l**, IPA ‘Diseases and Functions’ enriched in the DEG from classical monocytes cluster 1. The ones represented were selected among the ones with z-score > 2 or < −2, and p-value < 0.01. **m**, Monocyte subsets in the scRNAseq of PBMCs from human AD and HC donors (HC, *N* = 22; AD, *N* = 28) from Ramakrishnan et al.^23^. Unpaired t-test. **n**, DEG in AD CD14^+^ monocytes compared to HC CD14^+^ monocytes. Genes highlighted in light purple have log_2_FC > 0.25 or < −0.25 and p-value < 0.01, and those in dark purple are the top 10 DEG. **o**, Upstream regulators of CD14^+^ monocytes as calculated by IPA. Molecules highlighted in light purple have a z-score > 1 or < −1 and a p-value < 0.05. **p**, Venn Diagram showing overlapping of the upregulated IPA ‘Diseases and Functions’ enriched in the classical monocyte subcluster #1 and CD14^+^ human AD monocytes, selected among the ones with z-score > 1 or < −1, and p-value < 0.05.

We envisioned that blockade of CXCR4 would increase the circulation of monocytes in the blood. Administration of the CXCR4 antagonist, AMD3100, to WT and 5xFAD mice resulted in elevated levels of circulating monocytes (Extended Data Fig. 6a, b). It also mediated monocyte infiltration in the brain, since its blockade was linked with a partial reduction of the number of infiltrating myeloid cells in the brain and its borders (Extended Data Fig. 6c). Using *in vitro* transwell assays, we detected that monocytes derived from the AD migrate less toward CXCL12, the ligand for CXCR4, suggesting a dysfunctional state (Extended Data Fig. 6d, e). Altogether, this data highlights the role of CXCR4 in the migratory impairment of 5xFAD monocytes.

The BM harbors the differentiation of not only monocytes, but also all the myeloid cells. To better determine the extent of BM impairment in the AD models, other myeloid cell types were analyzed by flow and mass cytometry. No significant effects were observed in the abundance of neutrophils in the femur BM, the blood, or the spleen of 12-month-old 5xFAD when compared to age-matched WT (Extended Data Fig. 5f-h). The absence of effects on neutrophil numbers was also confirmed in the BM of the APP^NL-G-F^ model (Extended Data Fig. 5i). In contrast, dendritic cell (DC) differentiation was altered in the 5xFAD BM, showing a significant decrease in pre-DCs and an increase in type 1 conventional DCs (cDC1) (Extended Data Fig. 5j, k). These results suggest that the alterations in BM myelopoiesis in the AD models primarily affect monocytes and other related myeloid cells, such as DCs, but not the granulocytic lineage.

For a deeper characterization of the monocytic compartment in the AD BM, we performed single-cell transcriptomic analysis of CD11b^+^ myeloid cells and their cKit^+^ progenitors isolated from 12-month-old 5xFAD and age-matched WT. Using unsupervised clustering and UMAP, we detected a slight decrease in monocyte cluster abundance, which was accompanied by a significant decrease in non-classical monocytes (Fig. 3e-g, Extended Data Fig. 7a,b), validating the flow cytometry data (Fig. 3b, c). Further analysis of the classical monocytes in our single-cell transcriptomic dataset revealed several subpopulations (Fig. 3h), among which a specific cluster (Cluster 1) was increased in the BM of 5xFAD mice compared to WT animals (Fig. 3i). Analysis by IPA, revealed that DEG in this cluster were regulated by IFN-α (Fig. 3j, k), suggesting that IFN-I also affects the phenotype of mature classical monocytes. The DEG specific for this cluster were related to “Viral infection”, “Inflammatory response”, and “Adhesion of myeloid cells” (Fig. 3l), suggesting a viral-like inflammatory response in the BM of 5xFAD mice. Overall, our findings reveal an altered monocyte phenotype in the 5xFAD BM, characterized by heightened inflammatory activation, augmented accumulation of classical monocytes versus non-classical monocytes, and increased expression of CXCR4, which likely impairs their release into the circulation.

Analyzing the neutrophilic compartment in our BM scRNAseq dataset, revealed no major changes in the proportions of neutrophils and immature neutrophils (Fig. 3e, f, Extended Data Fig. 7b). DEG of clusters 1 and 2 in the 5xFAD neutrophils, showed an alternative activation and shift in their phenotype, mostly in the cluster 1 (Extended Data Fig. 7c, d). The DEG in the mature neutrophil cluster 1 again predicted the upregulation of “Viral response”, “Accumulation of ROS”, and “Cytotoxicity”; while there was a downregulation of the “Quantity of myeloid cells” and the “Inflammatory response”, potentially showing an attempt to restrain the inflammatory state under the influence of a viral-like response (Extended Data Fig. 7e).

We further tested whether the observed results in the BM of AD mice also appear in AD patients by taking advantage of a recently published scRNAseq dataset of PBMCs from AD patients and healthy controls (HC)^23^. In this study, the authors have shown that the phenotype of monocytes in AD patients is significantly affected, adopting a proinflammatory phenotype^23^. We re-clustered all the cells from the published dataset and identified the monocyte clusters (Extended Data Fig. 7f, g). We then subclustered the monocytes and identified the classical and non-classical monocytes based on common gene markers (Extended Data Fig. 7h, i). The classical monocytes (*CD14*^+^) showed a trend to expand in the AD patients compared to HC, whereas the non-classical monocytes (*FCGR3A*^+^*CD14*^-^) decreased (Fig. 3m), resembling the differences in blood and BM from the mouse models of AD (Fig. 3b, c, Extended Data Fig. 5a, e). The differential gene expression testing confirmed the proinflammatory phenotype by upregulation of *IL1B*, chemokines and *NFKB*-related genes (Fig. 3n). Interestingly, when analyzed by IPA, the DEG in these cells were predicted to be significantly regulated by *IFNα (family)*, *IFNβ (family)*, *IFNAR1*, *IRF5*, *STING1*, and poly I:C; all of them being involved in the IFN-I signaling pathway (Fig. 3o). The IPA functions related to the classical monocyte DEGs in the 5xFAD BM showed a significant overlap with those of the classical monocytes in the blood of AD patients (Fig. 3p). Among the overlapping “Diseases and Functions” from IPA, there were “Inflammatory response”, “Inflammation of CNS”, and “Activation of macrophages” as predicted to be upregulated (Extended Data Fig. 7j). Altogether, our findings show that the alteration of monocyte differentiation and proinflammatory phenotype are relevant to AD patients.

Monocyte and macrophage phenotypes in the 5xFAD central nervous system (CNS) have not been clearly characterized due to the low number of infiltrating cells and their predominant location at the brain borders (ie, choroid plexus, perivascular space, and meningeal layers). However, some studies with AD models provide histological evidence of their infiltration and crucial role in the brain parenchyma and its borders^15,44^. Furthermore, macrophages have been found in the brain parenchyma in postmortem sections from patients with advanced stages of AD^45^. Therefore, it is important to understand if the phenotype of peripheral monocytes in AD determines their function once they infiltrate into the brain or its borders. To address this question, we performed scRNAseq of the myeloid cells residing in the brain and brain borders (choroid plexus and leptomeninges). Unsupervised clusterization and annotation revealed that, besides the well-described acquisition of the DAM activation state by the microglia in the 5xFAD brain, there were no major changes in the proportion of other myeloid populations (Extended Data Fig. 8a, b). However, the DEG in the AD classical and non-classical monocytes revealed several genes related to neuroinflammation, damage of the vascular system, apoptosis, as well as downregulation of lipid metabolism (Extended Data Fig. 8c-f); all of them indicative of a pathological phenotype. When comparing these upregulated functions with those found in the mouse BM and the human PBMCs datasets, we found that more than half of them were conserved between compartments, showing a CNS correlation of those functions acquired in the periphery and at the origin of these cells (Extended Data Fig. 8g, h). These altered functions include “Encephalitis”, “Damage of vascular system”, and “Inflammation of absolute anatomical region”, among others (Extended Data Fig. 8i). Taken together, these results suggest that the pathological features detected in the periphery may drive a detrimental function of these cells within the CNS. The elevated levels of systemic circulating IFN-I detected in the plasma of 5xFAD mice (Fig. 2i), together with the transcriptomic analyses, led us to hypothesize that the observed effects on LT-HSC quiescence, impaired myelopoiesis, and pathological phenotype of mature monocytes might be linked and driven by circulating IFN-I.

### Bone marrow impairment in Alzheimer’s disease is rescued after blocking peripheral IFNAR1

To test the possible relationship between the impact of systemically elevated of IFN-I and the fate of the BM in the 5xFAD, we intraperitoneally treated these mice with an antagonistic monoclonal antibody against IFNAR1 for three weeks (Fig. 4a). Following the anti-IFNAR1 treatment, we detected an increase in the proportion of LT-HSC, and recovery of healthy quiescence in the HSPC compartment in the 5xFAD femur BM (Fig. 4b, c), reaching similar levels to those observed in WT (Fig. 1e). In addition, we tested the monocyte differentiation capacity of 5xFAD LSK cells *in vitro* following systemic anti-IFNAR1 treatment (Fig. 4d). We found that following IFNAR1 neutralization *in vivo*, there was a marked increase in the capacity of LSK differentiation to monocytes *in vitro*, indicating that blockade of IFNAR1 restored HSPC function (Fig. 4e). We also detected a significant increase in the numbers of CMP (Fig. 4f), and a significant reduction of CXCR4 expression on classical monocytes (Fig. 4g). No differences were detected in BM neutrophils in the anti-IFNAR1 treated 5xFAD mice (Extended Data Fig. 9a), showing a predominant effect of IFNAR1 activation in HSPC and monocytes in the 5xFAD BM.

**Fig. 4:**
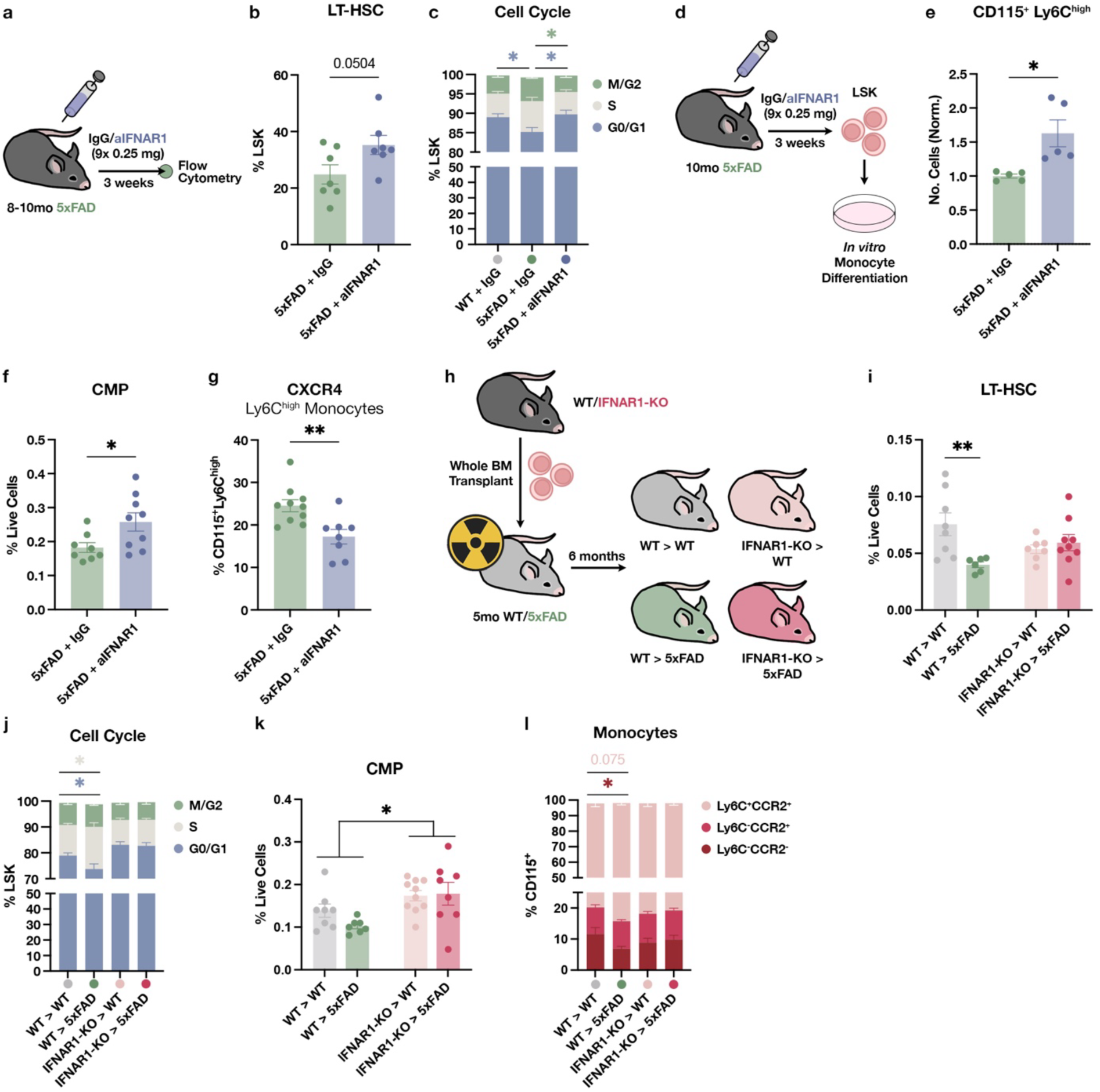
IFN-I blockade reverses pathological alterations in the 5xFAD BM. **a**, Peripheral aIFNAR1 injection scheme: 8-10-month-old AD mice were treated with monoclonal aIFNAR1 or IgG isotype nine times for 3 weeks; BM was then analyzed by flow cytometry. **b**, Abundance of LT-HSC in the 5xFAD BM treated with IgG or aIFNAR1 (5xFAD+IgG, *N* = 7; 5xFAD+aIFNAR1, *N* = 7). Unpaired t-test. **c**, Cell cycle analysis by flow cytometry of LSK from WT mice treated with IgG and 5xFAD mice treated with either IgG or aIFNAR1 (WT+IgG, *N* = 8; 5xFAD+IgG, *N* = 6; 5xFAD+aIFNAR1, *N* = 6). One-way ANOVA followed by Dunnett’s multiple comparisons test. **d**, AD animals were treated with IgG or aIFNAR1 for 3 weeks and then euthanized to isolate LSK from the BM. Sorted LSK were cultured for 1 week in the presence of monocyte differentiation factors, and the number of monocytes was measured by flow cytometry at the end of the culture. **e**, Absolute number of differentiated monocytes from LSK sorted from 5xFAD animals treated with IgG or aIFNAR1 after 7 days in culture. Values represent the mean of triplicates normalized with the average of the 5xFAD+IgG group. Data from two independent experiments. Unpaired t-test. **f**, Flow cytometry analysis of CMP abundance in the BM of 5xFAD animals treated with IgG or aIFNAR1 (5xFAD+IgG, *N* = 8; 5xFAD+aIFNAR1, *N* = 9). Unpaired t-test. **g**, Percentage of classical monocytes expressing CXCR4 in the BM of 5xFAD mice treated with IgG or aIFNAR1 (5xFAD+IgG, *N* = 9; 5xFAD+aIFNAR1, *N* = 7). Unpaired t-test. **h**, Total BM from WT or IFNAR1-KO mice was transplanted into 5-month-old irradiated WT or 5xFAD mice; animals were euthanized 6 months after the transplant, and the BM was analyzed by flow cytometry. **i**, Abundance of LT-HSC in WT and 5xFAD chimeric mice that received WT or IFNAR1-KO BM (*N* = 6-9 animals/group). Two-way ANOVA followed by Sidak’s multiple comparisons test. **j**, Cell cycle analysis of LSK by flow cytometry in the different chimeric groups analyzed (*N* = 8-10 animals/group). Unpaired t-test for each cell cycle phase comparing WT and 5xFAD receiving the same donor cell genotype. **k**, Abundance of CMP in WT and 5xFAD chimeric mice that received WT or IFNAR1-KO BM (*N* = 7-9 animals/group). Two-way ANOVA. **l**, Proportion of the different monocyte subsets analyzed by flow cytometry (*N* = 8-10 animals/group). Unpaired t-test for each monocyte subtype comparing WT and 5xFAD receiving the same donor cell genotype.

To further establish the role of IFN-I in driving these pathological features, we generated chimeric mice in which WT or 5xFAD recipients received total BM from either WT or IFNAR1-KO donors (Fig. 4h). To exclude a confounding contribution from skull BM hematopoiesis, animals were subjected to whole body irradiation prior to transplantation, reaching almost complete chimerism in peripheral blood (>90% on average, Extended Data Fig. 9b). To limit possible effects derived from irradiation-derived inflammation, BM transplants were performed when the animals were relatively young (5 month-old), and then the animals were tested 6 months later, when the disease was fully manifested (Fig. 4h). Analysis of the BM of IFNAR1-KO>5xFAD chimeras, showed that the LT-HSC levels were identical to those in the IFNAR1-KO>WT chimeric mice (Fig. 4i). Furthermore, the proportion of HSPC in G0/G1 phase in the BM of IFNAR1-KO>5xFAD chimeric mice was identical to that in the IFNAR1-KO>WT chimeras, in contrast to the significant reduction observed in the WT>5xFAD chimeric mice compared to WT>WT (Fig. 4j). In addition, there was an increased number of CMP in the BM of the animals that received IFNAR1-KO BM compared to those who received WT BM (Fig. 4k). Finally, the proportion of non-classical and classical monocytes in the BM of IFNAR1>5xFAD chimeras was not affected by the disease, in contrast to the reduction in the WT>5xFAD compared to WT>WT mice (Fig. 4l). Again, no differences were detected in the BM neutrophils after IFNAR1-KO transplantation (Extended Data Fig. 9c). Finding a similar phenotype in the WT>5xFAD compared to the 5xFAD non-transplanted mice further supported our findings that the observed effects are non-cell autonomous but driven by systemic factors, in particular, IFN-I.

### Blocking IFNAR1 in classical monocytes boosts resident myeloid cell function in the brain

To assess the potential impact of the IFN-I-dependent impairment of myelopoiesis on the disease manifestation. IFN-I is a crucial element that disrupts protective microglial functions in AD brains^6,46,47^. Moreover, whole-body knock-out of the *Ifnar1* gene is also protective in mouse models of AD^48^. We hypothesized that the observed restoration of monopoiesis upon IFN-I blockade might boost monocyte-derived macrophage homing into the brain with consequential protective effects in AD.

First, to determine the impact of the transplanted BM derived from IFNAR1 mice on the functional performance of the 5xFAD recipients, we assessed the cognitive performance of IFNAR1-KO>5xFAD chimeric mice using the novel object recognition (NOR) test (Fig. 5a). While WT>5xFAD mice showed cognitive impairment, calculated as the decrease in the discrimination index, IFNAR1-KO>5xFAD chimeras were protected (Fig. 5b). This protection against cognitive loss observed in the 5xFAD mice that received IFNAR1-KO BM was correlated with a significant increase in neuronal survival in the subiculum, as compared to WT>5xFAD (Fig. 5c, d); this brain region was previously reported as one of the main areas susceptible to neurodegeneration in this mouse model^49^. The cognitive and neuronal protection in the 5xFAD mice led us to analyze the underlying mechanism that connects IFN-I signaling in the BM with brain protection.

**Fig. 5:**
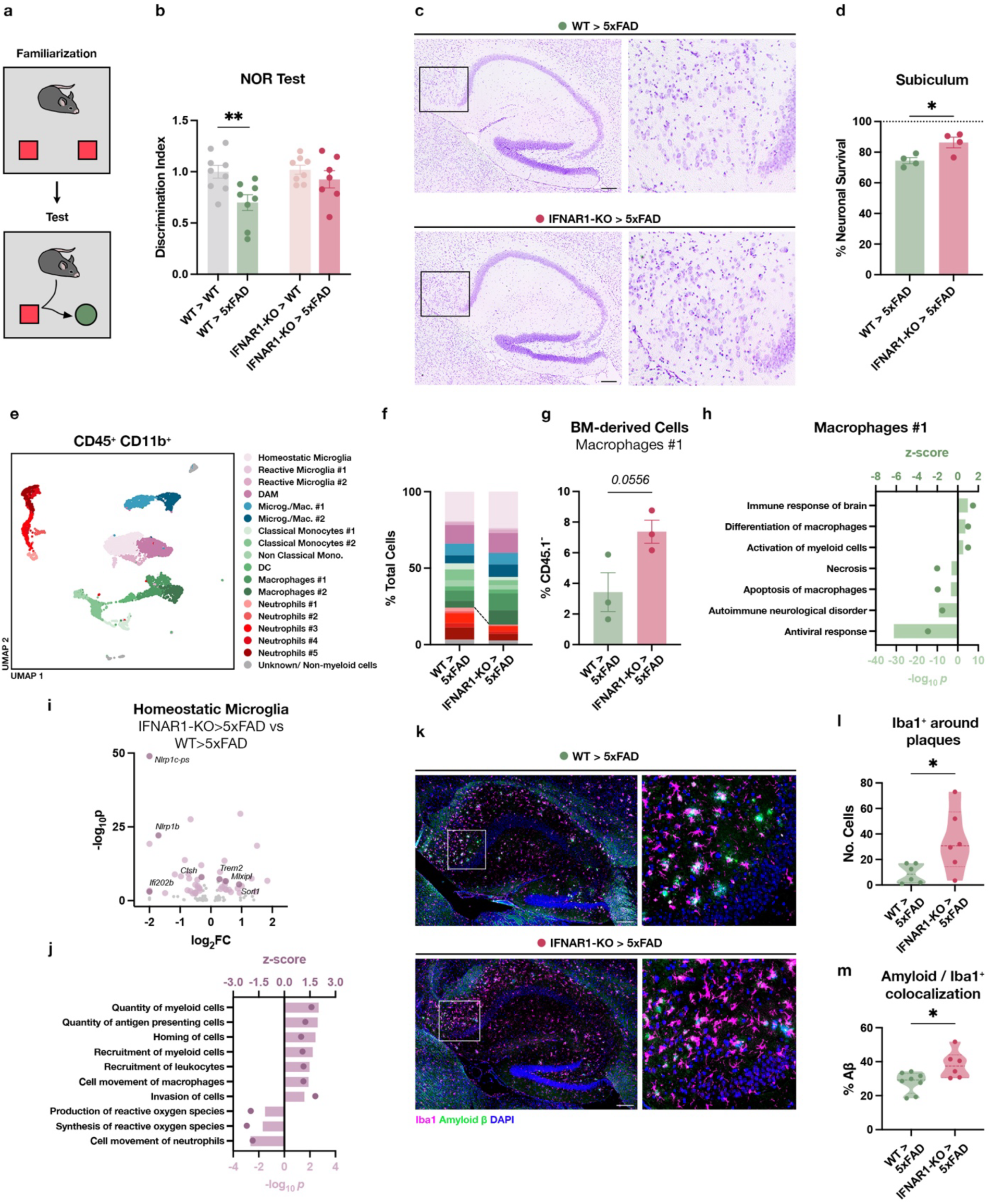
Immune peripheral knock-out of IFNAR1 remodels brain resident myeloid cell function and limits AD brain pathology. **a,** The novel object recognition (NOR) test was used to assess cognitive function. After habituation within the arena, mice were exposed to two identical objects in the familiarization phase. After 1-2 hours, one of the objects was replaced with a new one with a different shape, and the animals were exposed to them, measuring the time that they spent interacting with each object. **b**, Memory impairment is measured as a decrease in the discrimination index. The discrimination index was calculated as time spent with the novel object divided by the total time spent interacting with both objects. Data were normalized to the mean of the WT>WT mice measured on each testing day (*N* = 8-9 animals/group). Two-way ANOVA followed by Sidak’s multiple comparisons test. **c**, Representative pictures of cresyl violet staining of the hippocampus of WT>5xFAD and IFNAR1-KO>5xFAD chimeric mice. The subiculum is shown in the magnified inset. Scale bars represent 200 μm. **d**, Quantification of the neuronal survival in the 5xFAD chimeric animals as a percentage of the mean of neurons counted in the WT>WT animals (*N* = 4 animals/group). Unpaired t-test. **e**, UMAP showing clusters generated by scRNAseq analysis of CD11b^+^B220^-^TCRβ^-^ sorted from the brain of 11-month-old chimeric mice (WT>5xFAD, *N* = 4; IFNAR1-KO>5xFAD, *N* = 4; all were females). **f**, Proportions of the different myeloid clusters in the brain of WT>5xFAD and IFNAR1-KO>5xFAD animals. **g**, BM-derived cells (CD45.1^-^) contributing to the cluster of macrophages in the brain of chimeric 5xFAD mice. Unpaired t-test. **h**, IPA ‘Diseases and Functions’ enriched in the DEG from macrophages, selected among the ones with z-score > 1 or < −1, and p-value < 0.01. **i**, DEG in homeostatic microglia from IFNAR1-KO>5xFAD compared to WT>5xFAD. Genes highlighted in lavender have log_2_FC > 0.25 or < −0.25 and p-value < 0.01. **j**, IPA ‘Diseases and Functions’ enriched in the DEG from homeostatic microglia, selected among the ones with z-score > 1 or < −1, and p-value < 0.01. **k**, Representative pictures of Iba1 and amyloid beta (6E10) immunofluorescence of the hippocampus of WT>5xFAD and IFNAR1-KO>5xFAD chimeric mice. A region of the CA1 is shown in the magnified inset. Scale bars represent 200 μm. **l**, Quantification of the number of Iba1^+^ cells surrounding the amyloid plaques in the CA1 region of the hippocampus of WT>5xFAD and IFNAR1-KO>5xFAD chimeric mice (WT>5xFAD, *N* = 6; IFNAR1-KO>5xFAD, *N* = 6). Unpaired t-test. **m**, Quantification of the colocalization of amyloid beta and Iba1 staining in the CA1 region of the hippocampus of WT>5xFAD and IFNAR1-KO>5xFAD chimeric mice (WT>5xFAD, *N* = 8; IFNAR1-KO>5xFAD, *N* = 6). Unpaired t-test.

We analyzed by scRNAseq the profile of myeloid cells sorted from the brain of 5xFAD mice that received BM transplants from IFNAR1-KO or WT mice. Unsupervised clusterization and annotation of the clusters revealed a reduction in the number of infiltrating neutrophils (Fig. 5e, f, Extended Data Fig. 10a) and a proportional increase in the number of macrophages (Fig. 5e, f). This was validated by flow cytometry in a different batch of animals, despite a trend indicating a reduction in the global number of infiltrating myeloid cells (Extended Data Fig. 10b, c), there was an increased proportion of macrophages and a proportional decrease of neutrophils in the brains of IFNAR-KO>5xFAD mice compared to WT>5xFAD chimeras (Extended Data Fig. 10d). We further confirmed a similar profile of infiltrating myeloid cells following anti-IFNAR1 treatment in the 5xFAD mice (Extended Data Fig. 10e).

Infiltrating macrophages have shown the ability to exert protective roles in other models of CNS pathology^8–10^ and AD mouse models^17,50^. We hypothesize that blocking IFN-I signaling in the BM-derived cells was likely boosting the differentiation of peripheral monocytes into macrophages and affecting other resident myeloid cells in the 5xFAD brain. To dissect the origin of the different myeloid subsets in the brain, we took advantage of the CD45.1/CD45.2 mismatch between endogenous (CD45.1^+^CD45.2^+^) and transplant-derived cells (CD45.1^-^ CD45.2^+^). By flow cytometry analysis, we detected that more than 90% of the CD44^+^CD11b^+^ cells in the brain were CD45.1^-^, confirming their BM-transplant origin, whereas more than 70% of CD44^-^CD11b^+^ cells were CD45.1^+^. This validates our gating strategy and shows their long-lasting residency in the brain (Extended Data Fig. 10f, g). The origin of the different cell types was further validated in the scRNAseq dataset, employing the CITEseq technology and analyzing the protein expression of CD45.1 on the surface of the single-cells (Extended Data Fig. 10h). Based on protein values, CD45.1-positive or -negative cells were identified, confirming the endogenous origin of microglial clusters and the predominantly BM-transplant origin of the monocyte and neutrophil clusters (Extended Data Fig. 10i). The clusters identified as macrophages had more heterogeneous origin, probably comprising a mix of border-associated macrophages and monocyte-derived macrophages (Extended Data Fig. 10i). Interestingly, in the IFNAR1-KO>5xFAD mice, more macrophages were derived from the periphery, showing a boost in the infiltration and long-lasting residence of these cells (Fig. 5g). Since the majority of neutrophils, monocytes and some macrophages had a BM-transplant origin, their DEG in the brain of IFNAR1-KO>5xFAD mice were mostly related to IFN-I downregulation, as confirmed by IPA upstream regulator analysis (Extended Data Fig. 11a). The list of DEG analyzed by IPA showed increased activation of these cells, while there was a downregulation of “Necrosis” and “Autoimmune neurological disorder” (Fig. 5h). These cells also showed downregulation “Antiviral response”, likely resulting from the IFNAR1-KO genotype of the BM origin of some of these cells (Fig. 5h). This phenotype was similar to the one found in infiltrating classical monocytes, which showed also a marked downregulation of the “Inflammaroty response” (Extended Data Fig. 11b).

Resident cells were also affected by the nature of the infiltrating peripheral cells, namely IFNAR1-KO-derived or WT-derived. Homeostatic microglia and DAM showed several DEGs related to microglial function and DAM-phenotype acquisition (Fig. 5i, Extended Data Fig. 11c). In particular, some upregulated DEG in the homeostatic microglia from IFNAR1-KO chimeric 5xFAD mice were related to extracellular matrix remodeling (*Sorl1*, *Mmp2*), DAM-phenotype (*Trem2*), and lipid metabolism (*Mlxipl*); whereas some downregulated DEG were related to dampened inflammation (*Nlrp1b*, *Nlrp1b*-*ps*, *Ifi202b*, *Ctsh*) (Fig. 5i). IPA showed upregulated “Quantity of myeloid cells”, “Recruitment of myeloid cells” and downregulation of “Synthesis of reactive oxygen species”, among others (Fig. 5j). A similar phenotype was present in DAM, showing a conserved effect in different microglial states (Extended Data Fig. 11d). This suggests an enhanced reactivity to pathology of the microglia, with a dampened inflammatory phenotype.

This shift in microglial phenotype was likely due to the infiltration of IFNAR1-KO cells and the alteration of cell-to-cell communication networks between them. To detect differentially expressed cell-to-cell communication pathways, we ran the CellChat pipeline^51^. There were several upregulated and downregulated communication pathways between the classical monocytes and the resident myeloid cells (microglia and macrophages) (Extended Data Fig. 11e, f). Among the upregulated pairs of molecules in the IFNAR1-KO chimeric mice were the fibronectin-integrin pathway that can modulate microglia/macrophage polarization and phagocytosis^52,53^ (Extended Data Fig. 11e). On the other hand, among the downregulated pathways there were several described as detrimental in AD models, such as the ones related to complement 3 (C3)^54^, the immune checkpoint TIM-3 (encoded by *Havcr2*)^55^, or the amyloid precursor protein (APP) interaction with CD74 and TNFRSF21^56,57^ (Extended Data Fig. 11f). Altogether, our analysis suggests that infiltrating cells communicate with resident myeloid cells to orchestrate a benefitial phenotype in the brain of IFNAR1-KO>5xFAD chimeric mice.

Amyloid beta deposition is an early event in the 5xFAD mouse model and is a pathological hallmark of AD. Microglia, and particularly DAM, can compact and eliminate amyloid plaques^2^. We therefore explored whether modulating the phenotype of the infiltrating myeloid cells by affecting their differentiation in the IFNAR1-KO transplanted AD mice could improve microglial function. In the CA1 portion of the hippocampus, a region strongly related to AD cognitive decline, there were no changes in the size of plaques or their number between IFNAR1-KO and WT chimeric mice (Fig. 5k, Extended Data Fig. 11g, h). However, there was a significant increase in the number of Iba1^+^ cells surrounding these plaques (Fig. 5l), and a greater proportion of amyloid was colocalizing with these Iba1^+^ cells (Fig. 5m), suggesting an enhanced ability of the myeloid cells in the brain to sense and demarcate the amyloid plaques.

Overall, our results demonstrate that IFN-I signaling disrupts healthy monopoiesis in the 5xFAD bone marrow, thereby impairing macrophage infiltration into the AD brain. Targeting peripheral IFN-I signaling prevented this impairment, improved microglial phenotype, and halted cognitive decline in the 5xFAD mouse model of AD.

## Discussion

Monocyte-derived macrophages play a protective role in AD^14–16^. This suggests that peripheral monocytes may act as an intrinsic defense mechanism against neurodegeneration. However, the spontaneous recruitment of monocytes to the brain in AD appears to be inadequate. In this study, we demonstrate that the systemic environment in AD adversely affects monocyte function, impairing their capacity to support the brain in managing AD-related pathology.

Specifically, we showed in a mouse model of AD that hematopoiesis is disrupted, manifested by impairment of HSC quiescence and reduction of their abundance. We further found that this impairment limits myeloid differentiation, particularly the generation and maturation of new monocytes. Moreover, beyond the reduction in the myeloid progenitors, the produced monocytes display a pro-inflammatory phenotype and viral-like response. This phenotype was further validated by reanalyzing a published dataset of PBMCs from AD patients^23^. Here, we found that blocking IFN-I signaling, either by systemic administration of neutralizing monoclonal antibody or by creating chimeric mice in which the host BM was replaced with IFNAR1-KO BM, rescued normal HSC function, thereby reversing the BM impairments observed in AD mice. The peripheral blockade of IFN-I signaling also modulated infiltrating myeloid cells in the brain, reducing neutrophil infiltration, promoting monocyte/macrophage differentiation, and altering monocyte-microglia communication. Furthermore, when the IFN-I response was blocked in the peripheral immune cells, AD animals showed cognitive improvement and reduced neuronal loss compared to control AD mice.

The influence of CNS inflammation on hematopoiesis has been studied in the context of autoimmune diseases and injuries^19–22,30^. In these cases, the transmission of elevated levels of inflammatory mediators from the injured tissue to the BM is relatively rapid, triggering an activation of the hematopoiesis with a surge in new immune cells reaching the borders of the affected area in the brain or the spinal cord^19,30,31^. Our present study shows that chronic, progressive neurodegenerative diseases such as AD follow a different pattern. Instead of an increase in the myeloid bias of the BM of 5xFAD mice, we detected a marked decrease in the monopoietic potential.

A recent study has shown that the skull BM shows a particular enhancement and expansion during aging, increasing its capacity to generate new immune cells^32^. This privilege of the skull BM seems to be lost in AD, since it behaved similarly to the femur BM, exhibiting the same loss of LT-HSC and the same impairment in the monopoiesis *in vitro*. Of note, it has been recently demonstrated that the skull BM is connected to the dural meninges through microchannels that allow the direct delivery of myeloid cells^30,31^. While under acute injuries or infections, the skull and vertebrae BM respond rapidly, initiating emergency myelopoiesis mechanisms and sending myeloid cells to the CNS^8,19,20,30,31^, our results suggest that this might not be the case in chronic scenarios such as AD. Moreover, the fact that the removal of a peripheral reservoir of monocytes, such as the spleen, partially blocks monocyte recruitment to the AD brain^15^ suggests that monocytes, at least in part, need to go through circulation from the BM and the spleen before entering the AD brain.

BM hematopoiesis is greatly affected by the soluble proinflammatory factors that increase in aging, impacting the immune fitness of aged mice^58,59^. This age-dependent bias in immune cell differentiation limits the healing capacity of myeloid cells in the context of acute brain injury^60^. In the case of AD, the impairment of BM function is likely a result of the chronic exposure of the BM to brain-derived inflammatory signals. This aligns with our current findings, which show that impaired myelopoiesis initially occurs in the skull and later also in the femur BM niche. In fact, we are confirming previous reports of increased IFN-I production in the brain of AD mouse models^6,46^, which has been previously reported to negatively impact microglial activity in AD^6,46^ and during aging^61,62^. Although increased systemic levels of IFN-I were detected in the 5xFAD mice, further studies are needed to identify its source. Our study provides novel evidence that these deleterious effects of IFN-I extend beyond the brain in AD, and have a causal role in the impairment observed in BM function.

The role of IFN-I in the BM has been intensely studied as an important modulator of HSC health and differentiation^26,37–39,63^. The loss of LT-HSC and the HSPC entry into the activated cell cycle observed here in the 5xFAD BM is a well-defined effect of chronic IFN-I signaling^26,37^ but was never described before in the context of neurodegenerative diseases. We showed here that circulating IFN-I in AD mice impairs not only monopoiesis but also the transcriptomic program of mature monocytes. This maladaptive response in response to IFN-I was further detected in human CD14^+^ classical monocytes from AD patients, highlighting the translational potential of our findings. This maladaptive response to chronic peripheral IFN-I exposure could explain the limited spontaneous macrophage recruitment to resolve brain pathology in the context of AD. Furthermore, our findings are in line with recent reports showing that the epigenetic and transcriptomic program of circulating monocytes from AD patients exhibit a proinflammatory bias, more pronounced than that of any other immune cell type^23^. Moreover, research analyzing mature immune cells in the BM of AD mice suggests that these alterations are already evident during their differentiation process in the BM^64^. When combined with our findings, these results suggest that the circulating monocytes show an impaired phenotype acquired early in the differentiation process, preventing them from resolving AD brain pathology.

Previous reports have shown that BM transplantation from young animals is beneficial in mouse models of AD^64,65^. While Mishra et al. suggest that only WT BM is protective, Sun et al. demonstrated that even young BM from AD mice has a protective effect. Our findings indicate that the impairment of the BM is the result of a dysfunctional milieu that causes long-term stem cell impairment. This explains our observation that 5xFAD mice receiving WT BM continued to exhibit cognitive impairment. The only significant improvement was noted when 5xFAD mice received BM from IFNAR1-KO donors. These apparent discrepancies with previously reported results may stem from differences in the timing of the BM transplants and cognitive assessment. In the present study, BM transplantation was performed six months before cognitive assessment, and well before disease manifestation, providing sufficient time for the WT HSPC to be affected by the pathological signals present in the BM environment in the 5xFAD animals. Moreover, at the time of bone marrow transplantation, the host microenvironment had not yet undergone pathological changes. Therefore, the host BM and the transplanted WT BM were likely similar, with divergence emerging later in life. Another source of apparent discrepancies could be the manipulation of the donor cells. In the study from Mishra et al., only 5xFAD HSPC and not the WT HSPC were transfected with a reporter lentivirus, which could account for undetected anomalies in the 5xFAD HSPC triggered by anti-viral responses. We propose that the long-term signaling by IFN-I in the BM of 5xFAD mice impairs HSPC function and modifies monocyte phenotype, preventing a protective response by these cells in the AD brain.

Studies from several independent groups have shown that infiltration of monocytes/macrophages to the AD brain is protective^14–17^. Moreover, we and others have previously demonstrated that inhibition of the PD-L1/PD-1 axis arrests disease progression through a mechanism that is dependent on monocyte-derived macrophages ^50,66–68^. In addition, treatment with stimulating factors that boost myeloid differentiation and function, such as G-CSF and GM-CSF, is beneficial in mouse models of AD^69,70^ and is now being tested in clinical trials^71^. The present study emphasizes that the spontaneous reparative mechanism in AD could be impaired by IFN-I signaling in the BM, which blocks monocyte differentiation and skews the profile of these cells to a non-resolving phenotype. According to our findings, targeting chronic IFN-I signaling in the BM might boost the immune response in AD patients, and thus ameliorate brain pathology. Finding a balance between the preservation of a healthy anti-viral response and the suppression of the deleterious IFN-I response is needed to translate our findings to a clinical setting. Also, additional studies in HSPC from AD patients are needed to better understand if our findings are relevant to other forms of AD.

The current study supports the emerging understanding that AD is not only a disorder of the brain; rather, a significant component of the disease involves dysfunction of the systemic immune system^1,72,73^. Even if not the primary cause, this dysfunction acts as a significant catalyst for disease progression. Specifically, we present new evidence that immune system dysregulation, driven by IFN-I, is an integral component of AD pathology. Our findings also shed light on the connection between chronic anti-viral responses and the increased risk of developing AD^74,75^. Furthermore, they suggest a promising therapeutic avenue by targeting myelopoiesis to combat AD.

## Methods

### Animals

The heterozygous 5xFAD transgenic mice (on a mixed C57BL/6J-SJL background), which express familial AD mutant forms of human *APP* (the Swedish mutation, K670N/M671L; the Florida mutation, I716V; and the London mutation, V717I), and the mutant form of human *PSEN1* (M146L/L286V) transgenes under transcriptional control of the neuron-specific mouse Thy-1 promoter (5xFAD line Tg6799; The Jackson Laboratory) was used as an AD mouse model. Age-matched littermates that did not express the transgenes were used as WT controls.

Homozygous knock-in APP^NL-G-F^ transgenic mice (on C57BL/6J background), which express familial mutant forms of human *APP* (the Swedish mutation, KM670/671NL, the Iberian mutation, I716F, and the Arctic mutation, E693G) replacing the murine *App* gene, were used as an alternative model of familial AD and were provided by RIKEN Brain Science Institute (Japan). Age-matched C57BL/6J mice (Envigo) were used as controls.

UBC-GFP (JAX stock #004353, The Jackson Laboratory) and IFNAR1-KO (MMRRC stock #32045, The Jackson Laboratory) were used as BM donors or recipients, as indicated. C57BL/6J animals (Envigo) were used as WT controls for the generation of chimeric animals and for *in vitro* assays.

Males and females were used in all experiments in balanced proportions unless otherwise stated in the figure legend. Genotyping was performed by PCR analysis of ear-clip DNA. Mice were bred and maintained by the animal breeding center of the Weizmann Institute of Science and the animal facilities of the University of Navarra. All experiments detailed here were approved by the Institutional Animal Care and Use Committee (IACUC) of the Weizmann Institute of Science and by the Ethical Committee for Animal Testing at the University of Navarra.

### Flow Cytometry

Blood, spleen, and BM from the skull or the femur were collected for flow cytometric analyses. Blood was withdrawn from the cheek before perfusion. After perfusion with ice-cold PBS, skull BM was isolated by smashing the calvaria bones in a mortar after cleaning them from dural meninges and muscles. The femur and tibia BM were flushed out with FACS buffer using a syringe. The whole spleen was dissected after perfusion with ice-cold PBS and smashed. After erythrocyte lysis with RBC lysis buffer (BioLegend), samples were stained with the LIVE/DEAD Fixable Violet Kit (Invitrogen), and blocked with FcR-blocking Reagent (BioLegend). Cells were then incubated with fluorophore-conjugated antibodies (Extended Data Table 1) and acquired on a CytoFLEX S (Beckman Coulter) and a FACS Aria III (BD Biosciences).

For cell cycle analyses, the cells were fixed with PFA 4% (Thermo Fisher Scientific), washed with FACS buffer, and stained with DAPI (1/1000 in TX-100 0.1% in PBS) for 30 min at room temperature (RT). Cells were analyzed on a CytoFLEX S (Beckman Coulter) at a slow flow rate.

For analysis of brain immune cell infiltration, one hemisphere was collected after perfusion with ice-cold PBS for flow cytometric analyses. After cutting the brain tissue into small pieces, it was incubated with collagenase (Sigma-Aldrich) and DNase I (Roche) and gently homogenized to obtain a single-cell suspension. Myelin and cell debris were removed by centrifugation in 20% Percoll (Cytiva). Samples were then stained with the LIVE/DEAD Fixable Violet Kit (Invitrogen), and blocked with FcR-blocking Reagent (BioLegend). Afterward, cells were incubated with fluorophore-conjugated antibodies (Extended Data Table 1) and acquired on a CytoFLEX S (Beckman Coulter) and a FACS Aria III (BD Biosciences).

### Mass Cytometry

Femur BM was collected as described for flow cytometry analysis. After erythrocyte lysis, cells were incubated with anti-FcR 148Sm antibody (BD Biosciences, in-house conjugated as previously described^76^). Cells were barcoded with Cd-conjugated anti-mouse CD45 antibodies, incubated with anti-mouse CD34 APC (clone: SA376A4, 1:100, Biolegend). After washing, cells were incubated with a panel of metal-conjugated antibodies (Extended Data Table 2). After washing, cells were stained with Cell-ID Cisplatin and fixed in 4% formaldehyde. Fixed cells were acquired on a Helios CyTOF system (Fluidigm).

### Single-cell RNA sequencing

#### Single-cell RNA sequencing of HSPCs

For both LK and LSK populations, 3 x 10^4^ – 1 x 10^5^ cells were sorted after cell-multiplexing-oligo staining. Equal numbers of LK and LSK from each sample were combined at a 1:1 ratio, and loaded into two lanes of a 10X Chromium chip, following the manufacturer’s instructions for a final number of 33,000 cells loaded per lane. In total, seven femur samples were analyzed (four 5xFAD and three WT, 12-month-old). Library preparation was performed following ‘Chromium Next GEM Single Cell 3ʹ Reagent Kit v3.1 (Dual Index)’ instructions. Samples were sequenced in an Illumina Novaseq 6000, calculating an average of 25,000 reads per cell. Alignment was performed with CellRanger (v6.1.2). Downstream analyses were done using Seurat (v4.3) in R (v4.3.1). Quality control (QC) was performed, and cells with less than 100 or more than 5,000 detected genes were removed. A total of 25,946 cells were selected and normalized to 10,000 UMIs per cell and logarithmically transformed. Highly variable genes were selected using the “FindVariableFeatures” method. UMAP visualizations were obtained from 20 PCA components, and clusters were defined at a resolution of 0.5 using the K-nearest neighbor graph-based method. Cell types were annotated using typical marker genes for the different hematopoietic populations. One cluster of cells with high mitochondrial gene content was removed from the analysis. Differential gene expression was performed using the “FindMarkers” method using the Wilcoxon Rank Sum test. Pathway enrichment and upstream regulator analysis of differentially expressed genes were performed using Ingenuity Pathway Analysis (Qiagen).

#### Myeloid cell and HSPCs single-cell RNA sequencing

For mature myeloid cell single-cell transcriptomics, 4x10^5^ TER119^-^TCRβ^-^B220^-^CD11b^+^ cells and 5-9x10^4^ Lineage^-^cKit^+^ cells were sorted after cell-multiplexing-oligo staining. Before loading into the chip, CD11b^+^ and cKit^+^ from each sample were combined at a 7:3 ratio and loaded in two lanes of a 10X Chromium GEM-X chip, following the manufacturer’s instructions, for a final number of 2.9x10^4^ cells loaded per lane. In total, eight femur samples were loaded (four 5xFAD and four WT, 12-months-old). Samples were incubated with TotalSeq-B anti-mouse Hashtags (BioLegend) before sorting and multiplexing. Library preparation was done following the ‘Chromium GEM-X Single Cell 3ʹ Reagent Kit v4 (with Feature Barcodes)’ protocol. Samples were sequenced in an Illumina Novaseq X, calculating an average of 25,000 reads per cell. Alignment was performed with CellRanger (v6.1.2).

Downstream analyses were done using Seurat (v4.3) in R (v4.3.1). Quality control (QC) was performed, and cells with less than 100 or more than 6,000 detected genes were removed. 25,946 cells were selected and normalized to 10,000 UMIs per cell and logarithmically transformed. Highly variable genes were selected using the “FindVariableFeatures” method. UMAP visualizations were obtained from 20 PCA components, and clusters were defined at a resolution of 0.5 using the K-nearest neighbor graph-based method. Cell types were annotated using typical marker genes for the different hematopoietic populations. One cluster of cells with high mitochondrial gene content was removed from the analysis. Pathway enrichment and upstream regulator analysis of differentially expressed genes were performed using Ingenuity Pathway Analysis (Qiagen).

#### Myeloid cell single-cell RNA sequencing from the brain of untreated WT and 5xFAD mice

For brain myeloid cell single-cell transcriptomics, 4-9x10^4^ CD45^+^CD11b^+^ cells were sorted and fixed with the GEM-X Flex Sample Preparation v2 Kit following the manufacturer’s instructions for long-term storage. Cells were hybridized with mouse probes and barcodes for each mouse using the GEM-X Flex Sample Preparation v2 Kit. Cells were pooled for the washing and loaded in a 10X Chromium GEM-X Flex chip, following the manufacturer’s instructions, for a final number of 5.6x10^4^ cells loaded in one lane. In total, four samples were loaded (two 5xFAD and two WT, 12-months-old). Library preparation was done following the ‘Chromium GEM-X Flex Gene Expression v2’ protocol. Samples were sequenced in an Illumina Novaseq X, calculating an average of 10,000 reads per cell. Alignment was performed with CellRanger (v9.0.0). Downstream analyses were done using Seurat (v4.3) in R (v4.3.1). Quality control (QC) was performed, and cells with fewer than 200 detected genes, more than 4,500 detected genes, or more than 5% of mitochondrial genes were removed. 39,249 cells were selected and normalized to 10,000 UMIs per cell and logarithmically transformed. Highly variable genes were selected using the “FindVariableFeatures” method. UMAP visualizations were obtained from 25 PCA components, and clusters were defined at a resolution of 0.5 using the K-nearest neighbor graph-based method. Cell types were annotated using typical marker genes for the different hematopoietic populations. Pathway enrichment and upstream regulator analysis of differentially expressed genes were performed using Ingenuity Pathway Analysis (Qiagen).

#### Myeloid cell single-cell RNA sequencing from the brain of chimeric 5xFAD mice

Total brain cells were isolated as described above for flow cytometry analysis and incubated with CD45.1 TotalSeq-B Antibodies (BioLegend). After washing, 2.5-4.5x10^4^ CD11b^+^TCRβ ^-^B220^-^ cells were sorted and loaded in a 10X Chromium GEM-X OCM chip, following the manufacturer’s instructions, for a targeted cell recovery of 3,000 cells per sample. In total, eight samples were loaded (four WT>5xFAD and four IFNAR1-KO>5xFAD, 11-months-old). Library preparation was done following the ‘GEM-X Universal 3’ Gene Expression v4 4-plex (with Feature Barcodes)’ protocol with On-Chip Multiplexing. Samples were sequenced in an Illumina Novaseq X, calculating an average of 25,000 reads per cell for the gene expression library and 5,000 reads per cell for the feature barcode library. Alignment was performed with CellRanger (v9.0.1). Downstream analyses were done using Seurat (v4.3) in R (v4.3.1). Quality control (QC) was performed, and cells with fewer than 200 detected genes, more than 7,000 detected genes, or more than 5% of mitochondrial genes were removed. 6,980 cells were selected and normalized to 10,000 UMIs per cell and logarithmically transformed. Highly variable genes were selected using the “FindVariableFeatures” method. UMAP visualizations were obtained from 30 PCA components, and clusters were defined at a resolution of 0.5 using the K-nearest neighbor graph-based method. Cell types were annotated using typical marker genes for the different hematopoietic populations. Pathway enrichment and upstream regulator analysis of differentially expressed genes were performed using Ingenuity Pathway Analysis (Qiagen). Cell-to-cell communication networks were inferred with CellChat^51^ (v2).

### Bulk RNA sequencing from classical monocytes

For classical monocyte sequencing, single-cell BM suspensions from WT mice treated with PBS or poly I:C were prepared and stained with fluorochrome-conjugated antibodies, as above described. To this end, 1.5 x 10^5^ classical monocytes (CD11b^+^CD115^+^Ly6C^high^) were sorted in a FACS Aria III in purity mode and then resuspended in Qiagen Lysis buffer. The total RNA was extracted using the RNeasy Micro Kit (Qiagen), and cDNA libraries were prepared with the QuantSeq 3’ mRNA-Seq V2 Library Prep Kit with UDI (Lexogen). The libraries were sequenced in an Illumina Novaseq X, calculating an average of 30 million reads per sample. The reads were aligned using STAR (v 2.7.3a) to the mouse reference genome M24 (Gencode, GRCm38). A threshold of 0.5 CPM was set, and the differential gene expression was calculated with DESeq2 (v 1.36.0). Pathway enrichment and upstream regulator analysis of differentially expressed genes were performed using Ingenuity Pathway Analysis (Qiagen).

### Measurement of IFN-I by ELISA

Blood from 12-month-old WT and 5xFAD mice was collected in EDTA-coated tubes and centrifuged at 2,000 g to collect the plasma. Femur BM from 12-month-old WT and 5xFAD was collected by flushing the bones with FACS buffer and then centrifuged at 12,000 g to collect the supernatant. The brains from WT and age-matched 5xFAD were collected after perfusion, homogenized and and lysed in Lysis Buffer 2 (R&D Systems). For the detection of IFN-I, the Quantikine Mouse IFNα All Subtype ELISA Kit (R&D Systems) and Quantikine Mouse IFNβ ELISA Kit (R&D Systems) were used following the manufacturer’s instructions in duplicates. Data were acquired using an Infinite 200 PRO microplate reader (Tecan).

### Sorting and in vitro monocyte differentiation assay

BM single-cell suspension was prepared and stained with fluorochrome-conjugated antibodies as described above for flow cytometry analysis. HSPCs defined as Lineage^-^ Sca1^+^cKit^+^ (LSK) were sorted in flat-bottom 96-well plates and treated with SCF, M-CSF, and GM-CSF (all at 10 ng/mL, PeproTech) in IMDM supplemented with FBS (Gibco), glutamine, beta-mercaptoethanol, penicillin, and streptomycin. Media and growth factors were partially refreshed after 5 days in culture. Murine IFN-α2 (eBioscience) or human IFN-β (PeproTech) was added to the media at the indicated doses from the initiation of the cell culture, and refreshed together with the other factors. After 7 days in culture, cells were harvested, and the number of differentiated monocytes was analyzed by flow cytometry based on CD11b, CD115, and Ly6C expression.

### Chimeric mouse generation

For the generation of whole-body chimeras, 4-5-month-old WT or 5xFAD mice were irradiated in an X-Rad320 X-ray irradiator with a full dose of 800 cGy. The following day, total BM from femurs and tibias of C57BL/6J or IFNAR-KO mice was flushed, and the erythrocytes were lysed. A total of 3 x10^6^ BM cells were injected (iv) into the irradiated recipients. The animals received enrofloxacin in their drinking water for three weeks. The percentage of chimerism was assessed six weeks after the transplant based on CD45.2/CD45.1 and IFNAR expression by flow cytometry analysis of the blood of the recipient animals.

### Competitive bone marrow transplantation

BM cells were isolated from 12-month-old WT or 5xFAD mice, and single-cell suspensions were prepared and stained with fluorochrome-conjugated antibodies as described above for flow cytometry analysis. From each animal, 6 x 10^3^ LSK were sorted and injected together with 1.2 x10^6^ total BM competitor cells from UBC-GFP mice into irradiated UBC-GFP recipient animals. The animals were irradiated on day −1, as described above. Chimerism and myeloid cell production by the transplanted LSK (GFP^-^ CD45.1^+^ CD45.2^+^) were evaluated in the peripheral blood by flow cytometry every four weeks starting four weeks after the transplantation.

### Polyinosinic:polycytidylic acid treatment

For experiments using polyinosinic:polycytidylic acid (poly I:C), C57BL6/J animals (8 weeks old, Envigo) or age-matched IFNAR1-KO were used. Mice were treated with poly I:C (ip, 10 mg/kg, Sigma-Aldrich) every 48 hours for one week. Controls received PBS. Animals were euthanized 24 hours after the last dose.

### CXCR4 blockade

For *in* vivo blockade of CXCR4, AMD3100 (MedChem Express) was dissolved in EtOH and then in corn oil (10%/90%) and injected daily (ip, 5 mg/kg, 6 doses) to either WT or 5xFAD mice. Mice were euthanized 24h after the last dose.

### Peripheral IFNAR1 blockade

InVivoMAb anti-mouse IFNAR1 (BioXCell) or InVivoMAb mouse IgG1 isotype control (BioXCell) were injected (ip, 0.25 mg/mouse, 9 doses) every 48-72h to either WT or 5xFAD mice. Mice were euthanized 72 hours after the last dose.

### In vitro migration assay

BM monocytes were sorted using CD115-Microbeads (Miltenyi Biotec) and plated in supplemented RPMI in the upper chamber of a Transwell Plate (5.0 µm Pore Polycarbonate Membrane, Corning). Migration media containing murine CXCL12 (Peprotech) at the indicated concentrations were added to the lower chamber. Three hours after plating the cells, the plates were briefly centrifuged, and the migrated monocytes were measured by flow cytometry.

### Behavioral test

Mice were handled daily for at least seven days before the initiation of NOR test. The NOR test spanned two days and included three trials: habituation trial, day 1 (20 min session in the empty arena); familiarization trial, day 2 (10 min session in the arena with two identical objects located in opposite corners of the floor, approximately 9 cm from the walls on each side); test trial, day 2, 60-70 min after familiarization (6 min session in the arena with one of the objects replaced by a novel one). Familiar and novel objects were visually and tactilely distinct. Two identical arenas placed side by side were used simultaneously (one mouse in each arena). Each arena consisted of a 41.5×41.5 cm gray plastic box. Novel object preference was measured as a discrimination index, expressed as the time of interaction with the novel object relative to the time of interaction with both objects (%) during the test trial. A discrimination index above 50% indicates novelty recognition, with 50% indicating no recognition. After each trial, the arenas and equipment were wiped with 10% ethanol. Female and male mice were tested on different days. Time spent interacting with each object was measured by a blinded experimenter.

### Histological Analysis

After perfusion with ice-cold PBS, one brain hemisphere was fixed in 4% PFA overnight at 4°C. After that, samples were transferred to 1% PFA, embedded in paraffin, and sectioned at a thickness of 6 μm. Cresyl violet staining was performed in four consecutive sagittal sections to detect neuronal nuclei. Images were acquired in a Leica DMi8 microscope and analyzed with ImageJ (v2.14.0) by automated cell counting.

Immunofluorescence was performed by staining sagittal sections overnight with rabbit anti-Iba1 (1:150, Wako) and mouse anti-amyloid beta (clone: 6E10, 1:150, Biolegend), followed by secondary antibodies Cy3 goat anti-rabbit (1:300, Jackson ImmunoResearch) and Cy2 goat anti-mouse (1:300, Jackson ImmunoResearch). Sections were counterstained with DAPI (1:10,000, Sigma-Aldrich). Images were acquired in a Leica DMi8 microscope and analyzed with ImageJ (v2.14.0) by automated counting of the plaques. Iba1^+^ cells were counted in the area of the plaque and the 15 μm surrounding each plaque.

### Reanalysis of public datasets

Publicly available single-cell RNA-sequencing data from human blood (GSE226602) was analysed using the R (v4.3.3) package Seurat (v5.0.3). Raw data were subjected to quality control, filtered to retain those with >200 features, <20,000 UMIs, and <10% mitochondrial gene expression. Potential doublets were identified and removed using the scDblFinder package (v1.16). Data were normalised and scaled using SCTransform, and patient-derived batch effects were corrected using Harmony (v1.2). The corrected data were used for dimensionality reduction and visualisation with UMAP. Monocyte populations were identified by the expression of canonical markers (CD14, FCGR3A, MS4A7), isolated, and sub-clustered to define classical and non-classical subtypes. Differential gene expression analysis between Alzheimer’s Disease and healthy control monocytes was performed using a MAST (v1.2) test, with a significance threshold of an adjusted p-value < 0.05 and an absolute log2 fold-change ≥ 0.5. Pathway enrichment and upstream regulator analysis of differentially expressed genes were performed using Ingenuity Pathway Analysis (Qiagen).

### Statistical tests

The normal distribution of the data was checked by the D’Agostino-Pearson normality test. Normally distributed data was analyzed by unpaired t-test, one-way ANOVA, and two-way ANOVA. One-way ANOVA was followed by Dunnett’s multiple comparisons test, and two-way ANOVA was followed by Sidak’s multiple comparisons test. Data that didn’t follow a normal distribution was analyzed with the Mann-Whitney test. Statistical outlier detection was performed with the ROUT method.

## Data availability

Transcriptomic sequencing data generated in this study will be publicly available in Gene Expression Omnibus database upon acceptance of this manuscript. Complete lists of DEG and IPA results can be found in Supplementary Information. All other raw data are available from the corresponding authors upon request.

## Acknowledgments

The authors thank Merav Kedmi from the Life Sciences Core Facilities, the team from the Flow Cytometry Unit, the team from the Animal Facility at the Weizmann Institute of Science, and Ksenia Glonty from the Weizmann Institute of Science, for their technical support. Also, the authors thank Prof. Ido Amit and Prof. Gideon Schreiber for providing the GFP and IFNAR1-KO transgenic mouse strains. We also thank all members of the Schwartz laboratory for helpful discussions.

This study was supported by the Advanced European Research Council (grant no. 7417), the Israel Science Foundation (ISF research grant no. 991/16), and the ISF Legacy Heritage Bio-Medical Science Partnership (research grant no. 1354/15). We thank the Thompson Foundation for their generous support of our AD research. M.A.A. was supported by a postdoctoral fellowship from ‘Fundacion Ramon Areces’.

## Author contributions

M.A.A., L.B., and M.P. designed, performed, and interpreted the experiments and wrote the manuscript. M.S. and A.D. supervised the participants, secured funding for the study, and revised and wrote the manuscript. C.B. and M.K. performed experiments and revised the manuscript. G.C., S.P.C., J.M.P.R., S.M., A.I., P.A., Y.A., H.P., B.N., and M.E. assisted in the experiments. M.C.T and A.G.O. assisted with the design of some experiments and revised the manuscript. T.M.S. assisted in the design and supervised the mass cytometry experiments.

R.V. and J.J. assisted in the analysis of the RNAseq experiments. L.C. performed the genotyping of the mice.

---

**Extended Data Table 1:** Conjugated antibodies used in flow cytometry analysis and fluorescence-activated cell sorting.

| Antigen | Fluorophore | Clone | Supplier | Ref. No. | Dilution |
| --- | --- | --- | --- | --- | --- |
| B220 | BV421 | RA3-6B2 | BioLegend | 103251 | 1/200 |
| CCR2 | FITC | Y15-488.rMAb | BD Biosciences | 568414 | 1/100 |
| CD115 | PE | AFS98 | BioLegend | 135506 | 1/200 |
| CD117 (cKit) | PE | 2B8 | BioLegend | 105808 | 1/200 |
| CD117 (cKit) | BV605 | 2B8 | BioLegend | 105847 | 1/200 |
| CD11b | PE-Cy7 | M1/70 | BioLegend | 101216 | 1/200 |
| CD11b | BV605 | M1/70 | BioLegend | 101257 | 1/200 |
| CD11b | FITC | M1/70 | BioLegend | 101206 | 1/500 |
| CD135 | APC | A2F10 | BioLegend | 135309 | 1/100 |
| CD150 | PE-Cy5 | TC15-12F12.2 | BioLegend | 115912 | 1/200 |
| CD16/32 | BV421 | 2.4G2 | BD Biosciences | 752948 | 1/200 |
| CD31 | APC | 390 | BioLegend | 102410 | 1/200 |
| CD34 | PE-Dazzle | SA376A4 | BioLegend | 152210 | 1/100 |
| CD4 | PE-Cy5 | RM4-5 | BioLegend | 100513 | 1/200 |
| CD44 | FITC | IM7 | BioLegend | 103006 | 1/200 |
| CD45 | BV421 | 30-F11 | BioLegend | 103134 | 1/200 |
| CD45.1 | PE | A20 | BioLegend | 110708 | 1/200 |
| CD45.1 | APC | A20 | BioLegend | 110714 | 1/200 |
| CD45.2 | AF700 | 104 | BioLegend | 109822 | 1/200 |
| CD45.2 | FITC | 104 | BioLegend | 109806 | 1/200 |
| CD48 | APC-Cy7 | HM48-1 | BioLegend | 103432 | 1/200 |
| CD8 | AF700 | 53-6.7 | BioLegend | 100729 | 1/200 |
| CX3CR1 | PE | SA011F11 | BioLegend | 149006 | 1/200 |
| CXCR4 | BV421 | L276F12 | BioLegend | 146511 | 1/200 |
| Lineage Cocktail | Biotin | --- | Miltenyi Biotec | 130-092-613 | 1/20 |
| Ly-6C | AF700 | HK1.4 | BioLegend | 128024 | 1/100 |
| Ly-6C | PE-Dazzle | HK1.4 | BioLegend | 128044 | 1/200 |
| Ly-6C | APC | HK1.4 | BioLegend | 128016 | 1/200 |
| Ly-6C | BV711 | HK1.4 | BioLegend | 128037 | 1/200 |
| Ly-6G | BV605 | 1A8 | BioLegend | 127639 | 1/200 |
| Ly-6G | PE | 1A8 | BioLegend | 127608 | 1/500 |
| Ly-6A/E (Sca-1) | PE-Cy7 | D7 | BioLegend | 108114 | 1/200 |
| Streptavidin | BV650 | --- | BioLegend | 405232 | 1/200 |
| Streptavidin | FITC | --- | BioLegend | 405202 | 1/200 |
| Streptavidin | PE | --- | BioLegend | 405204 | 1/200 |
| TCRbeta | APC-Cy7 | H57-597 | BioLegend | 109220 | 1/200 |
| TER-119 | APC | TER-119 | BioLegend | 116212 | 1/200 |

**Extended Data Table 2:** Conjugated antibodies used in mass cytometry analysis.

| Antigen | Metal | Clone | Supplier | Dilution | Notes |
| --- | --- | --- | --- | --- | --- |
| CD117 cKit | 166Er | 2B8 | Fluidigm | 1/100 |  |
| APC | 162Dy | APC003 | Fluidigm | 1/100 |  |
| Sca-1 | 169Tm | D7 | Fluidigm | 1/100 |  |
| CD150 | 167Er | TC15-12F12.2 | Fluidigm | 1/100 |  |
| CD127 | 175Lu | A7R34 | Fluidigm | 1/100 |  |
| CD135 | 157Gd | A2F10.1 | BD Biosciences | 1/100 | In-house labeled |
| CD11c | 209Bi | N418 | Fluidigm | 1/100 |  |
| MHC-II | 174Yb | M5/114.15.2 | Fluidigm | 1/100 |  |
| CD123 | 147Sm | 5B11 | BioLegend | 1/100 | In-house labeled |
| pDCA1 | 161Dy | 927 | BioLegend | 1/100 | In-house labeled |
| XCR1 | 152Sm | ZET | Fluidigm | 1/100 |  |
| CD172a | 142Nd | P84 | BioLegend | 1/100 | In-house labeled |
| CX3CR1 | 164Dy | SA011F11 | Fluidigm | 1/100 |  |
| CD115 | 144Nd | AFS98 | Fluidigm | 1/100 |  |
| CD24 | 150Nd | M1/69 | Fluidigm | 1/100 |  |
| B220 | 176Yb | RA3-6B2 | Fluidigm | 1/100 |  |
| CD19 | 149Sm | 6D5 | Fluidigm | 1/100 |  |
| TCRbeta | 143Nd | H57-597 | Fluidigm | 1/100 |  |
| CD4 | 145Nd | RM4-5 | Fluidigm | 1/100 |  |
| CD8a | 168Er | 53-6.7 | Fluidigm | 1/100 |  |
| Ly-6G | 141Pr | 1A8 | Fluidigm | 1/100 |  |
| Ly-6C | 198Pt | HK1.4 | Fluidigm | 1/100 |  |
| CD11b | 196Pt | M1/70 | Fluidigm | 1/100 |  |
| Ter119 | 154Sm | Ter119 | Fluidigm | 1/100 |  |

**Extended Data Fig. 1:**
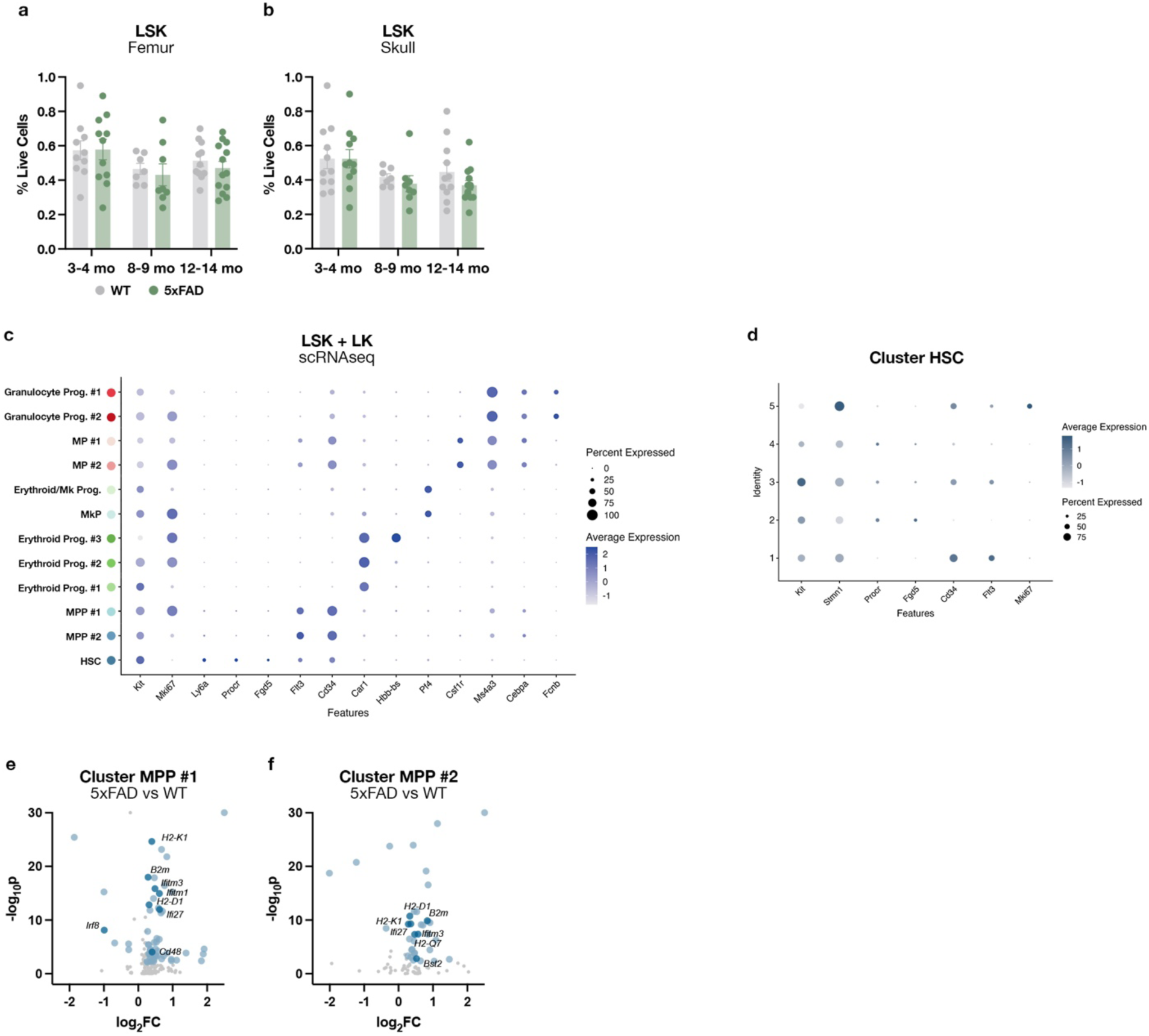
Hematopoietic stem cell quiescence in the skull bone marrow is lost in the 5xFAD mouse model of familial AD. **a,** LSK abundance in the femur BM of WT and 5xFAD animals at different time points analyzed by flow cytometry. Unpaired t-test. **b,** LSK abundance in the skull BM of WT and 5xFAD animals at different time points analyzed by flow cytometry. Unpaired t-test. **c,** Dot plot showing gene markers used for the general annotation of LSK and LK cells analyzed by scRNAseq from the femur BM of 12-14 month-old WT and 5xFAD mice (WT, *N* = 3; 5xFAD, *N* = 4; all were females). **d,** Dot plot showing gene markers used in the annotation of subclusters from the HSC cluster analyzed by scRNAseq from the femur BM of 12-14-month-old WT and 5xFAD mice. **e**, DEG of 5xFAD MPP cluster #1 compared to WT cells. Genes highlighted in light blue have log_2_FC > 0.25 or < −0.25, and p-value < 0.01, and those in dark blue are related to IFN-I signaling and MPP commitment. **f**, DEG of 5xFAD MPP cluster #2 compared to WT cells. Genes highlighted in light blue have log_2_FC > 0.25 or < −0.25, and p-value < 0.01, and those in dark blue are related to IFN-I signaling.

**Extended Data Fig. 2:**
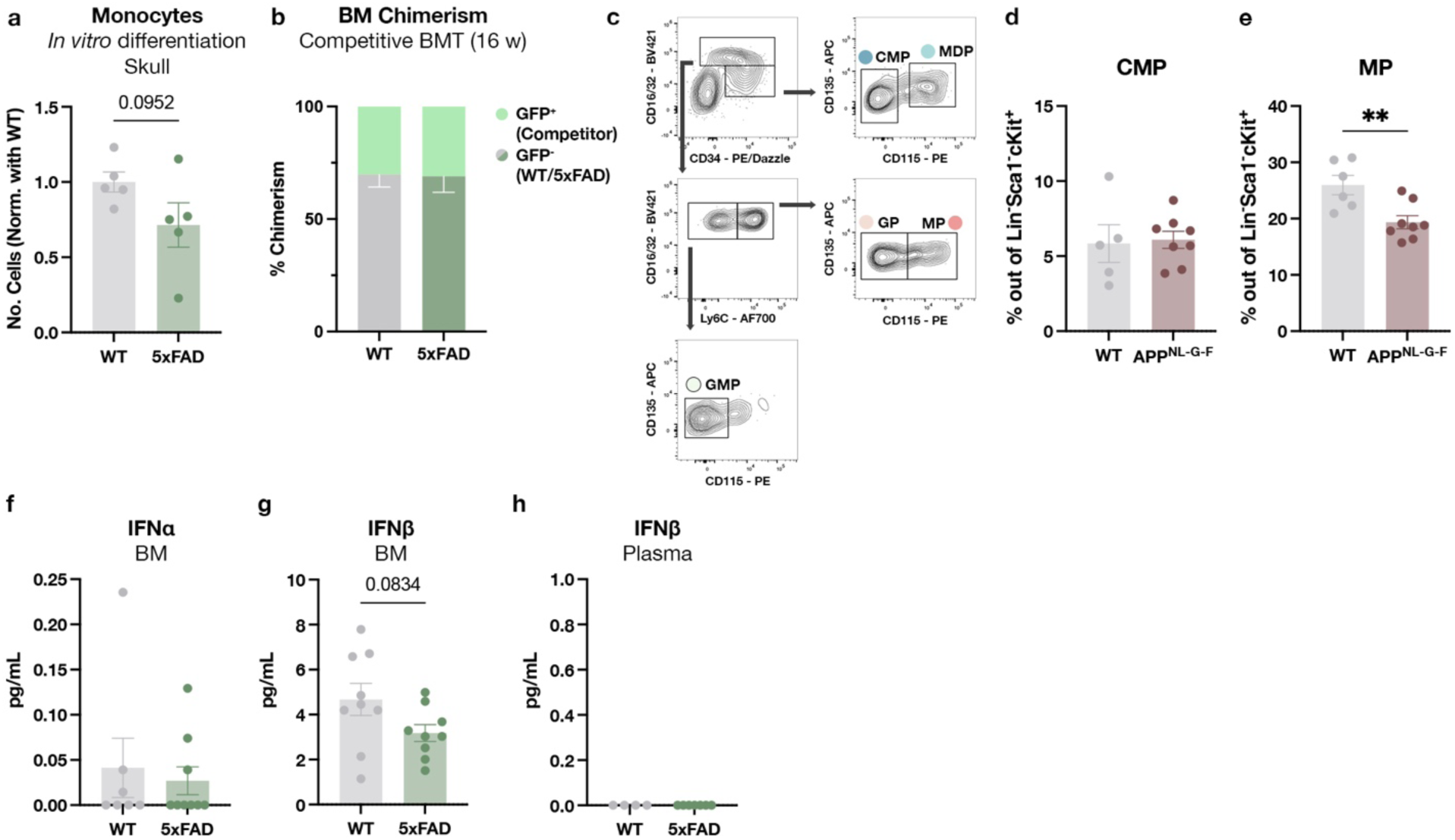
Myeloid progenitor differentiation is affected by neurodegeneration in the 5xFAD model and the APP knock-in model. **a**, Absolute number of differentiated monocytes from 12-month-old WT or 5xFAD LSK from the skull BM after 7 days in culture. Values represent the mean of triplicates normalized with the average of the WT group. Data from two independent experiments. Mann-Whitney test. **b**, Degree of chimerism in the BM of GFP animals 16 weeks after the transplant of WT or 5xFAD LSK and competitor GFP cells (WT, *N* = 7; 5xFAD, *N* = 9). **c**, Gating strategy for myeloid progenitor subset identification by flow cytometry. **d**, Flow cytometry analysis of CMP in the BM of 20-month-old APP^NL-G-F^ mice and age-matched WT (WT, *N* = 5; APP^NL-G-F^, *N* = 8). Unpaired t-test. **e**, Flow cytometry analysis of MP in the BM of 20-month-old APP^NL-G-F^ mice and age-matched WT (WT, *N* = 6; APP^NL-G-F^, *N* = 8). Unpaired t-test. **f**, IFNα concentration in the femur BM collected from 12-month-old WT and 5xFAD animals as analyzed by ELISA (WT, *N* = 7; 5xFAD, *N* = 9). **g**, IFNβ concentration in the femur BM collected from 12-month-old WT and 5xFAD animals as analyzed by ELISA (WT, *N* = 9; 5xFAD, *N* = 9). **h**, IFNβ concentration in the plasma collected from 12-month-old WT and 5xFAD animals as analyzed by ELISA (WT, *N* = 4; 5xFAD, *N* = 7).

**Extended Data Fig. 3:**
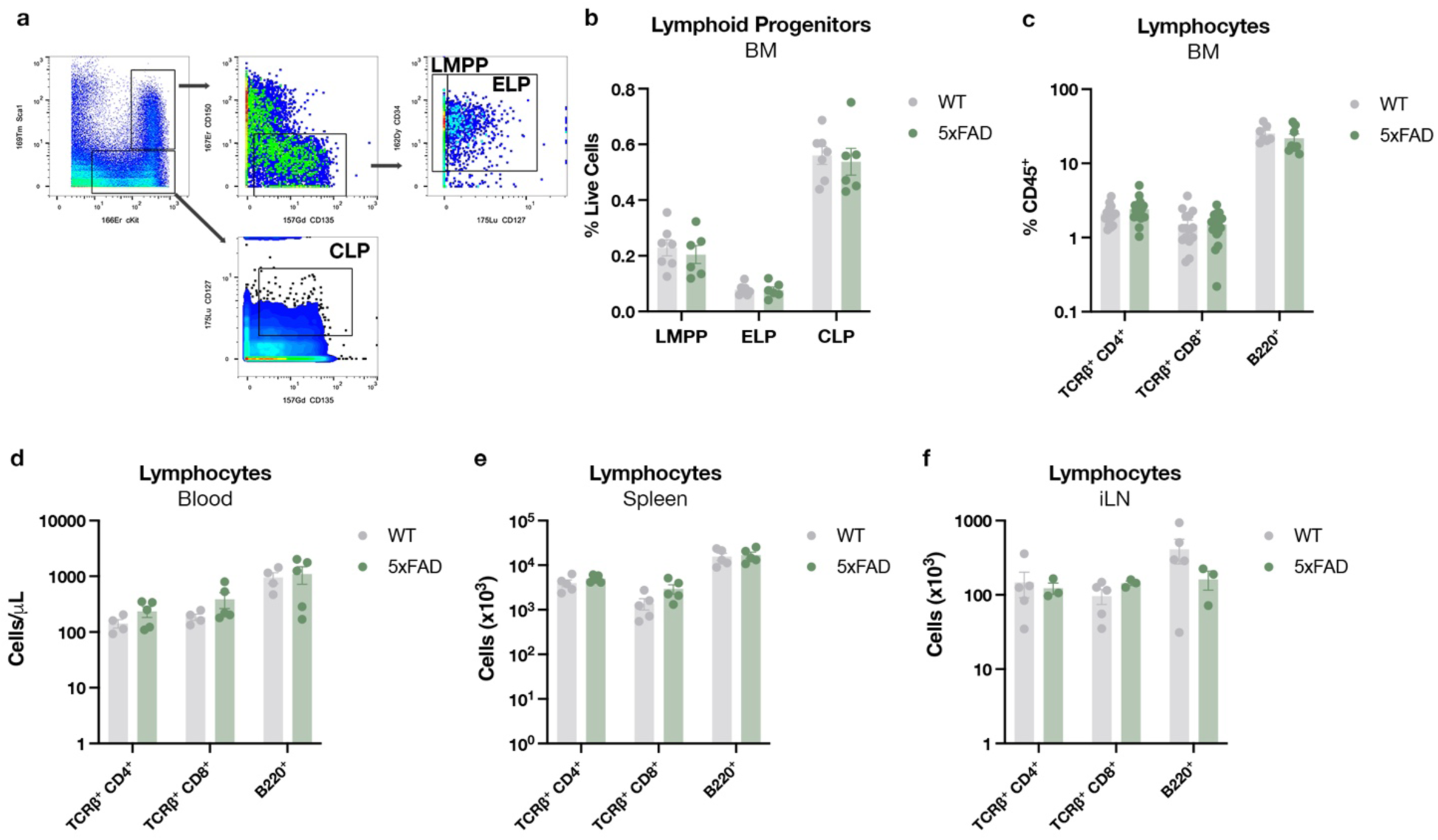
No evident alterations in the early lymphoid progenitors or lymphocytes in the 5xFAD femur BM. **a**, Gating strategy for lymphoid progenitor analysis by mass cytometry. **b**, Mass cytometry analysis of lymphoid multipotent progenitors (LMPP), early lymphoid progenitors (ELP), and common lymphoid progenitors (CLP) in the femur BM of 12-month-old WT and 5xFAD littermates (WT, *N* = 7; 5xFAD, *N* = 6). **c**, Flow cytometry analysis of T and B lymphocytes in the femur BM of 12-month-old WT and 5xFAD littermates (*N* = 7-17 animals per group). **d**, Flow cytometry analysis of T and B lymphocytes circulating in the blood of 12-month-old WT and 5xFAD littermates (WT, *N* = 4; 5xFAD, *N* = 5). **e**, Flow cytometry analysis of T and B lymphocytes in the spleen of 12-month-old WT and 5xFAD littermates (WT, *N* = 5; 5xFAD, *N* = 5). **f**, Flow cytometry analysis of T and B lymphocytes in the inguinal lymph node (iLN) of 12-month-old WT and 5xFAD littermates (WT, *N* = 5; 5xFAD, *N* = 3).

**Extended Data Fig. 4:**
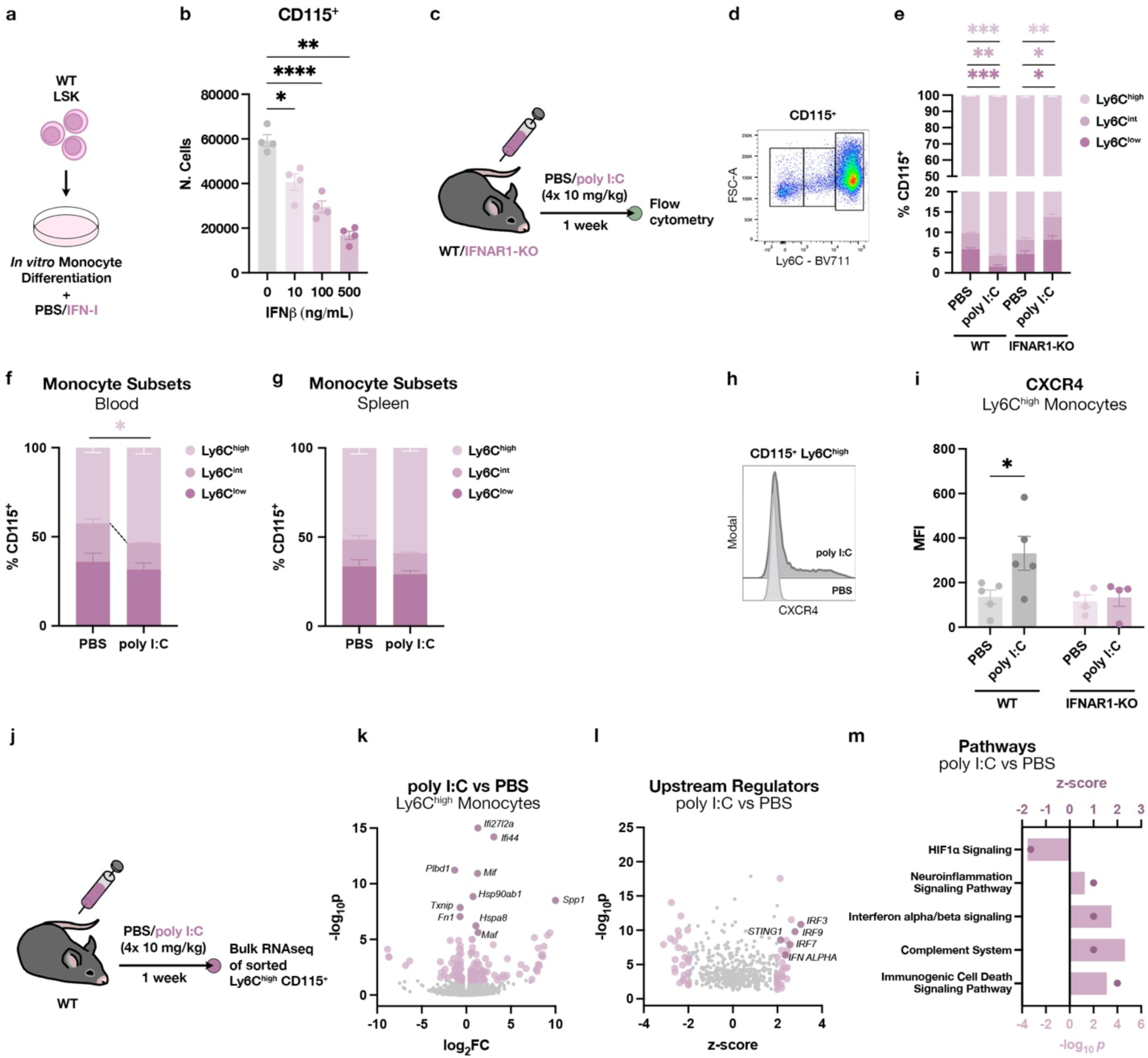
IFN-I signaling in the BM impacts the mature monocyte compartment. **a**, Sorted LSK were cultured for one week in the presence of IFNβ and monocyte differentiation factors. **b**, Absolute numbers of monocytes after one week *in vitro* following exposure to different doses of IFNβ. Repeated measures one-way ANOVA followed by Dunnett’s multiple comparisons test. **c**, WT or IFNAR1-KO were treated for one week with poly I:C, and then the BM was analyzed by flow cytometry. **d**, Gating strategy for monocyte subset analysis by flow cytometry. **e**, Monocyte subsets in the BM of PBS- or poly I:C-treated WT or IFNAR1-KO mice (*N* = 5/group). Unpaired t-test between PBS- vs poly I:C-treated of each genotype. **f**, Monocyte subsets in the peripheral blood of PBS- or poly I:C-treated WT mice (PBS, *N* = 5; poly I:C, *N* = 4). Unpaired t-test in each monocyte subtype between conditions. **g**, Monocyte subsets in the spleen of PBS- or poly I:C-treated WT mice (PBS, *N* = 5; poly I:C, *N* = 4). Unpaired t-test in each monocyte subtype between conditions. **h**, Histogram showing CXCR4 expression in classical monocytes in PBS- and poly I:C-treated WT mice. **i**, Median fluorescence intensity (MFI) of CXCR4 expressed by classical monocytes in the BM of PBS- or poly I:C-treated WT or IFNAR1-KO mice (*N* = 4-5/group). Unpaired t-test between PBS- vs. poly I:C-treated mice of each genotype. **j**, WT mice were treated with PBS or poly I:C for one week, and classical monocytes were sorted for bulk RNAseq analysis. **k,** DEG in classical monocytes sorted from poly I:C-treated animals versus controls. Genes highlighted in lavender have log_2_FC > 0.5 or < −0.5 and p-value < 0.05, and those in violet are the top-10 DEG. **l,** Upstream regulators as calculated by IPA. Molecules highlighted in lavender have a z-score > 2 or < −2 and p-value < 0.05. **m,** IPA ‘Pathways’ enriched in monocytes from pIC-treated animals, selected among the ones with z-score > 1 or < −1, and p-value < 0.05.

**Extended Data Fig. 5:**
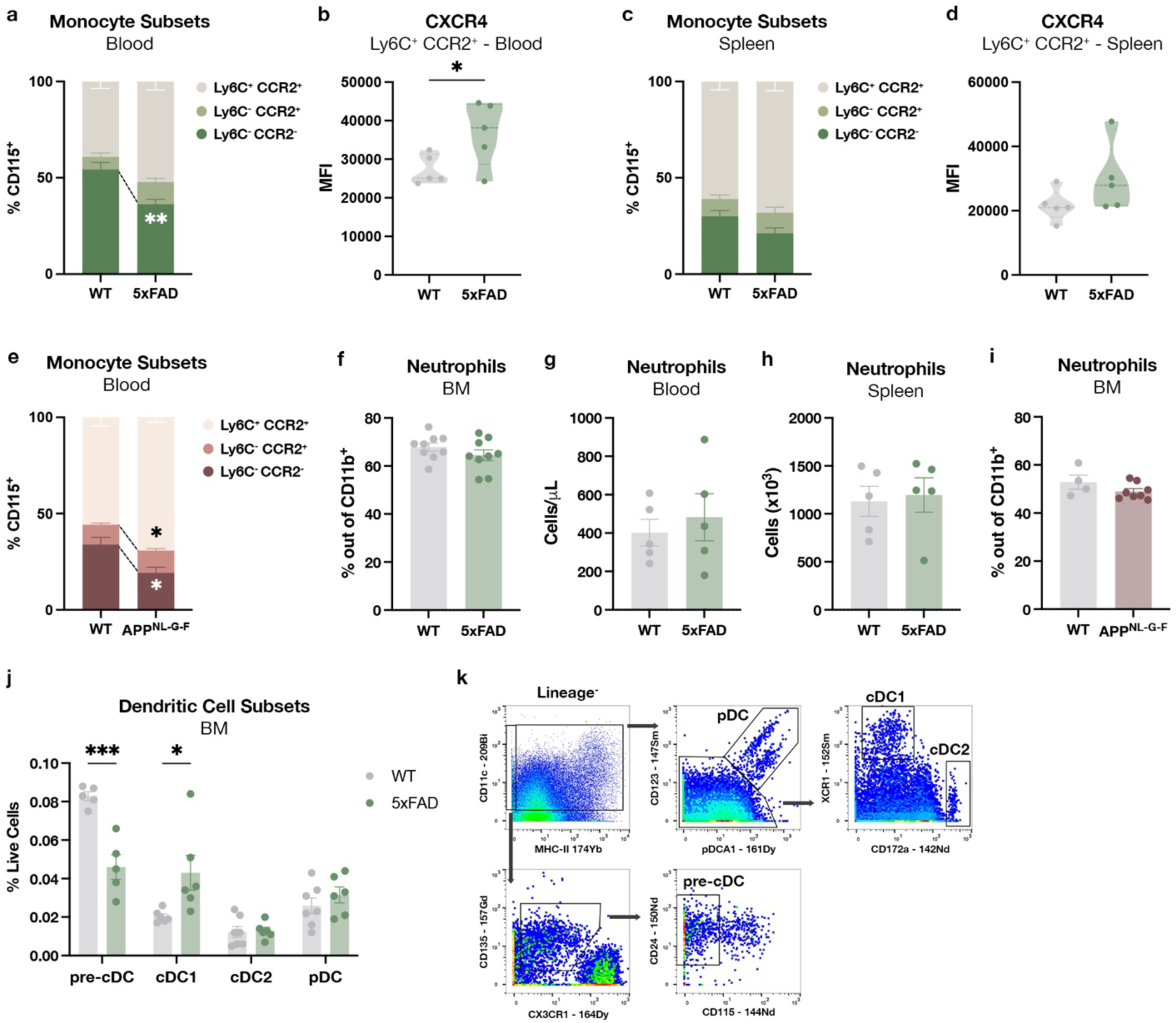
Circulating monocytes in the blood of AD mice exhibit phenotypic changes that are preserved from the BM phenotype. **a**, Monocyte subsets in the blood of 12-month-old WT and 5xFAD mice (WT, *N* = 5; 5xFAD, *N* = 4). Unpaired t-test for each subtype comparing WT and 5xFAD. **b**, Median fluorescence intensity (MFI) of CXCR4 expressed by classical monocytes in the blood of 12-month-old WT and 5xFAD mice (WT, *N* = 5; 5xFAD, *N* = 5). Unpaired t-test. **c**, Monocyte subsets in the spleen of 12-month-old WT and 5xFAD mice (WT, *N* = 5; 5xFAD, *N* = 5). Unpaired t-test for each subtype comparing WT and 5xFAD. **d**, Median fluorescence intensity (MFI) of CXCR4 expressed by classical monocytes in the spleen of 12-month-old WT and 5xFAD mice (WT, *N* = 5; 5xFAD, *N* = 5). Unpaired t-test. **e**, Monocyte subsets in the blood of 20-month-old WT and APP^NL-G-F^ mice (WT, *N* = 4; APP^NL-G-F^, *N* = 6). Unpaired t-test for each subtype comparing WT and APP^NL-G-F^. **f**, Flow cytometry analysis of neutrophil abundance in the BM of 12-month-old WT and 5xFAD mice (WT, *N* = 9; 5xFAD, *N* = 9). Unpaired t-test. **g**, Flow cytometry analysis of neutrophil abundance in the blood of 12-month-old WT and 5xFAD mice (WT, *N* = 5; 5xFAD, *N* = 5). Unpaired t-test. **h**, Flow cytometry analysis of neutrophil abundance in the spleen of 12-month-old WT and 5xFAD mice (WT, *N* = 5; 5xFAD, *N* = 5). Unpaired t-test. **i**, Flow cytometry analysis of neutrophil abundance in the BM of 20-month-old WT and APP^NL-G-F^ mice (WT, *N* = 4; APP^NL-G-F^, *N* = 8). Unpaired t-test. **j**, Mass cytometry analysis of dendritic cell subsets in the femur BM of 12-month-old WT and 5xFAD mice (*N* = 5-7 animals/group). Unpaired t-test. **k**, Gating strategy for dendritic cell subset analysis by mass cytometry.

**Extended Data Fig. 6:**
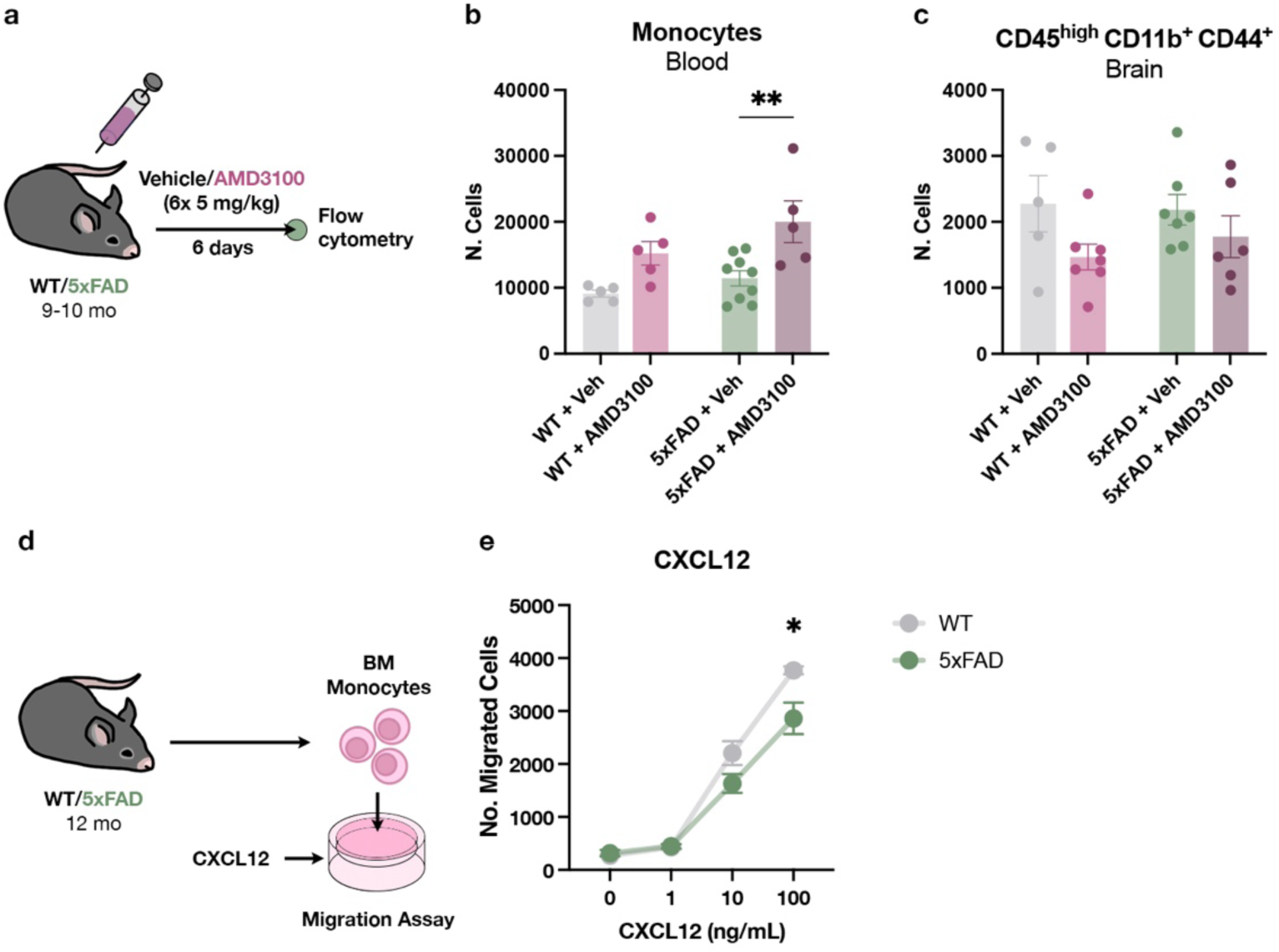
Monocyte migration is dependent on CXCR4, and the migratory ability of monocytes towards CXCL12 is impaired in the 5xFAD BM. **a**, Peripheral AMD3100 injection scheme: 9-10-month-old WT and 5xFAD mice were treated with AMD3100 or vehicle daily for six days; blood and brain were then analyzed by flow cytometry one day after the last dose. **b**, Number of circulating monocytes in the blood of WT or 5xFAD mice treated with AMD3100 or vehicle (*N* = 5-9 animals/group). Two-way ANOVA followed by Sidak’s multiple comparisons test. **c**, Number of infiltrating monocytes in the brain of WT or 5xFAD mice treated with AMD3100 or vehicle (*N* = 5-9 animals/group). Two-way ANOVA followed by Sidak’s multiple comparisons test. **d**, Monocytes from the BM of 12-month-old WT or 5xFAD were isolated and plated on top of a Transwell plate that was loaded in the lower chamber with different concentrations of CXCL12. The absolute number of migrated monocytes was measured 3 hours later in the lower chamber by flow cytometry. **e**, Number of migrated monocytes following increasing concentrations of CXCL12. Unpaired t-test for each concentration.

**Extended Data Fig. 7:**
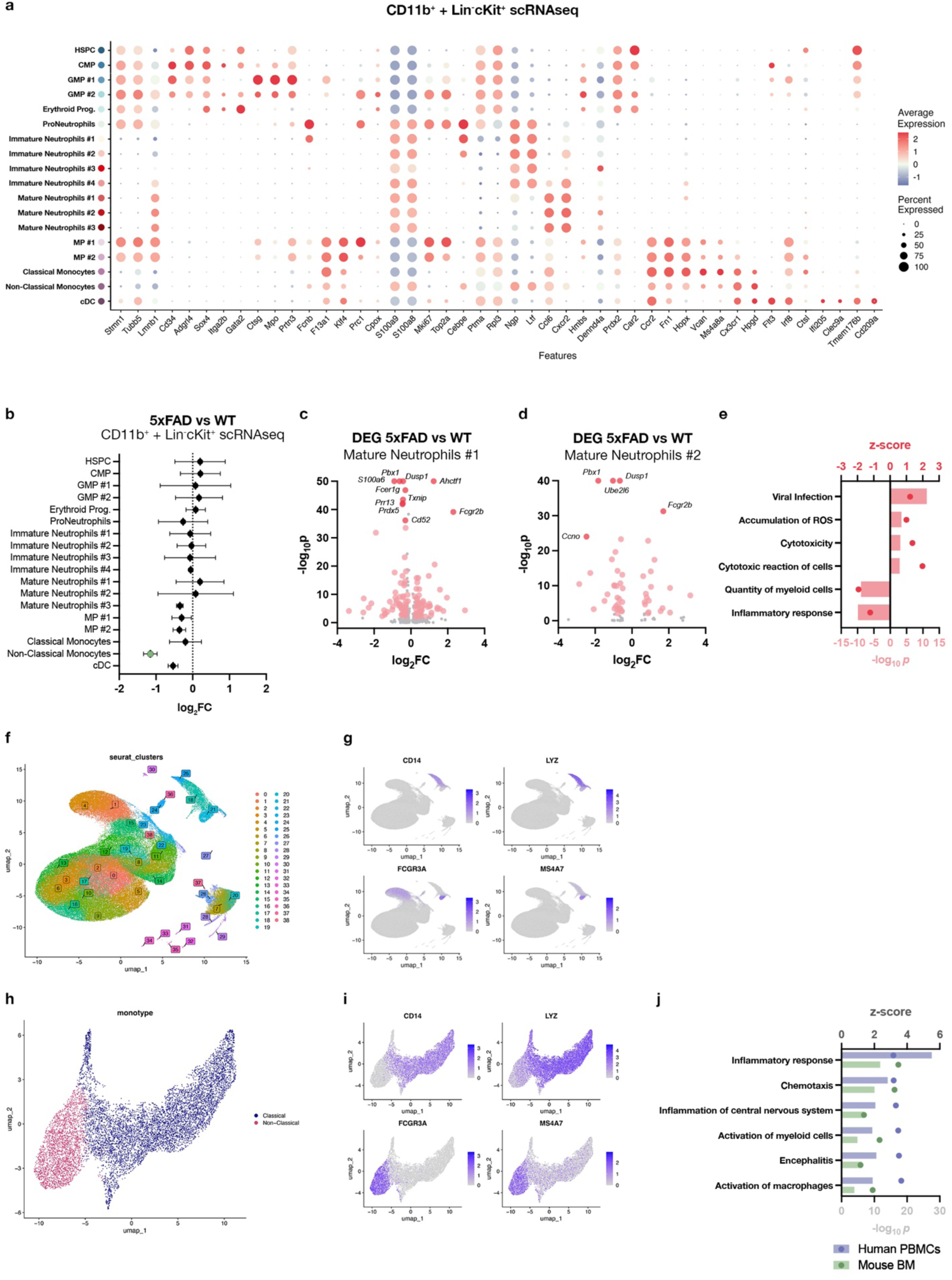
IFN-I signaling in the BM alters the mature myeloid compartment. **a,** Dot plot showing gene markers used for the general annotation of CD11b^+^ and LK cells analyzed by scRNAseq from the femur BM of 12-month-old WT and 5xFAD mice (WT, *N* = 4; 5xFAD, *N* = 4; all were females). **b,** Fold change of the proportion of general clusters from CD11b^+^ and Lin^-^cKit^+^ cells analyzed by scRNAseq. Values shown represent the mean ± SEM. **c**, DEG in mature neutrophils cluster #1 comparing 5xFAD to WT animals. Genes highlighted in pink have log_2_FC > 0.25 or < −0.25, and p-value < 0.01, and those in red are the top 10 DEG. **d**, DEG in mature neutrophils cluster #2 comparing 5xFAD to WT mice. Genes highlighted in pink have log_2_FC > 0.25 or < −0.25, and p-value < 0.01, and those in red are the top 5 DEG. **e,** IPA ‘Diseases and Functions’ predicted from the DEG from neutrophil cluster #1, selected among the ones with z-score > 1 or < −1, and p-value < 0.05. **f**, UMAP showing the re-clusterization of the human AD scRNAseq dataset from Ramakrishnan et al.^23^. **g**, Feature plot with gene markers of human monocytes. **h**, Subclusterization and annotation of the monocyte clusters from the human AD scRNAseq dataset. **i**, Feature plot with gene markers of human monocyte subsets. **j**, ‘Diseases and Functions’ shared between the 5xFAD mouse BM and the human AD PBMCs datasets, selected among the ones with z-score > 1, and p-value < 0.05.

**Extended Data Fig. 8:**
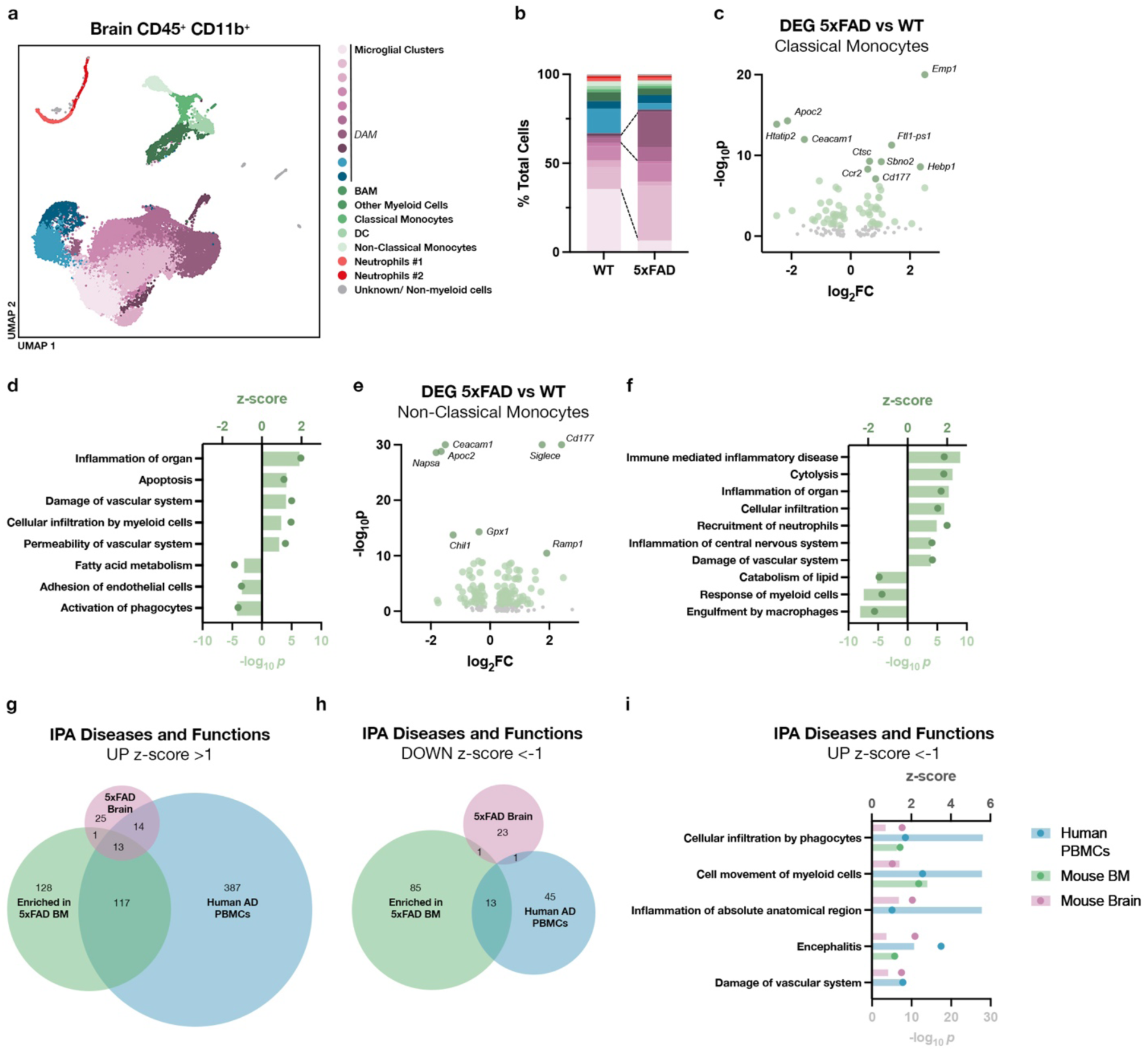
Infiltrating myeloid cells in the 5xFAD brain share features with those in the BM at the transcriptomic level. **a**, UMAP showing clusters generated by scRNAseq analysis of CD45^+^CD11b^+^ cells sorted from the brain of 12-month-old WT or 5xFAD mice (WT, *N* = 2; 5xFAD, *N* = 2; all were females). **b**, Proportions of the different clusters in the brain of WT and 5xFAD animals. **c**, DEG in classical monocytes comparing 5xFAD to WT mice. Genes highlighted in light green have log_2_FC > 0.25 or < −0.25, and p- value < 0.05, and those in dark green are the top 10 DEG. **d**, IPA ‘Diseases and Functions’ predicted from the DEG of 5xFAD classical monocytes. The ones represented were selected among the ones with z-score > 1 or < −1, and p-value < 0.01. **e**, DEG in non-classical monocytes comparing 5xFAD to WT mice. Genes highlighted in light green have log_2_FC > 0.25 or < −0.25, and p-value < 0.05, and those in dark green are the top 10 DEG. **f**, IPA ‘Diseases and Functions’ predicted from the DEG of the 5xFAD non-classical monocytes. The ones represented were selected among the ones with z-score > 1 or < −1, and p-value < 0.01. **g**, Venn diagram showing the overlapping IPA ‘Diseases and Functions’ increasing in the AD classical monocytes from the scRNAseq datasets generated from the 5xFAD mouse BM and brain, and the human AD PBMCs. **h**, Venn diagram showing the overlapping IPA ‘Diseases and Functions’ decreasing in the AD classical monocytes from the scRNAseq datasets generated from the 5xFAD mouse BM and brain, and the human AD PBMCs. **i**, IPA ‘Diseases and Functions’ predicted to increase in the classical monocytes from 5xFAD BM, brain, and the human AD PBMCs.

**Extended Data Fig. 9:**
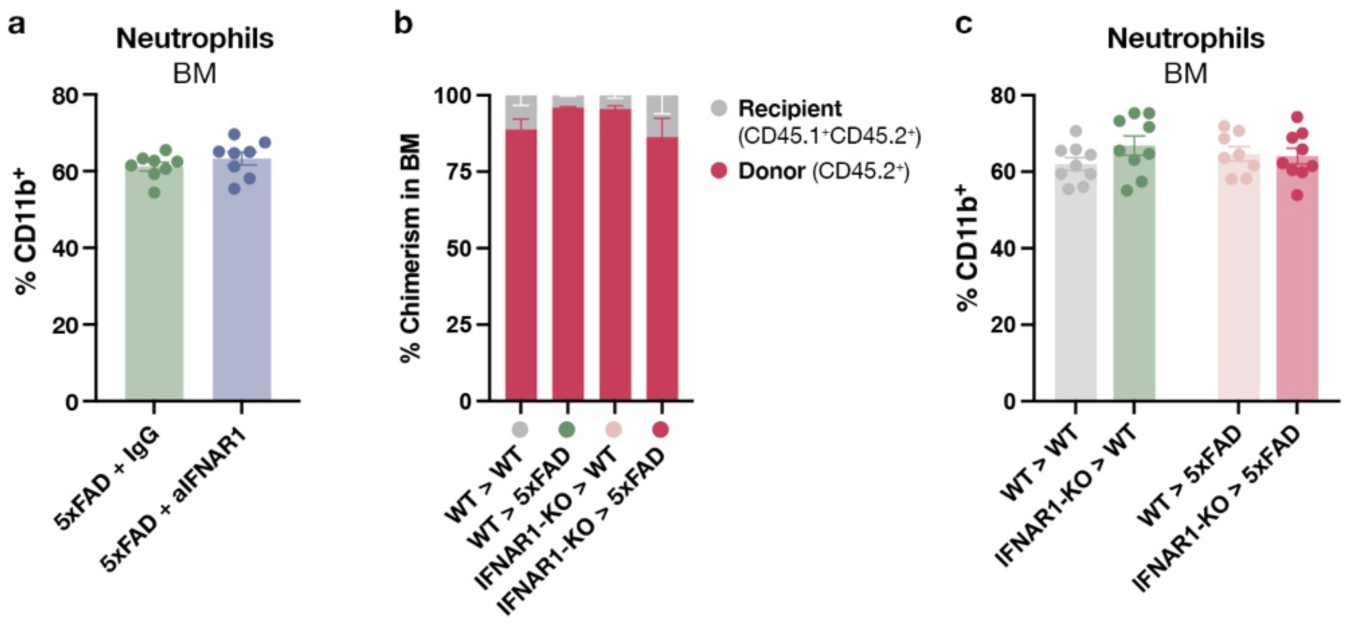
IFN-I signaling blockade does not alter neutrophil abundance in the BM. **a**, Flow cytometry analysis of neutrophils in the femur BM of 5xFAD mice after IgG or aIFNAR1 treatment (*N* = 8 animals/group). **b**, Chimerism in the BM of WT and 5xFAD mice that received total BM transplant from WT or IFNAR1-KO mice (*N* = 8-9 animals/group). **c**, Flow cytometry analysis of neutrophils in the femur of WT or 5xFAD chimeric mice that received WT or IFNAR1-KO BM (*N* = 8-9 animals/group).

**Extended Data Fig. 10:**
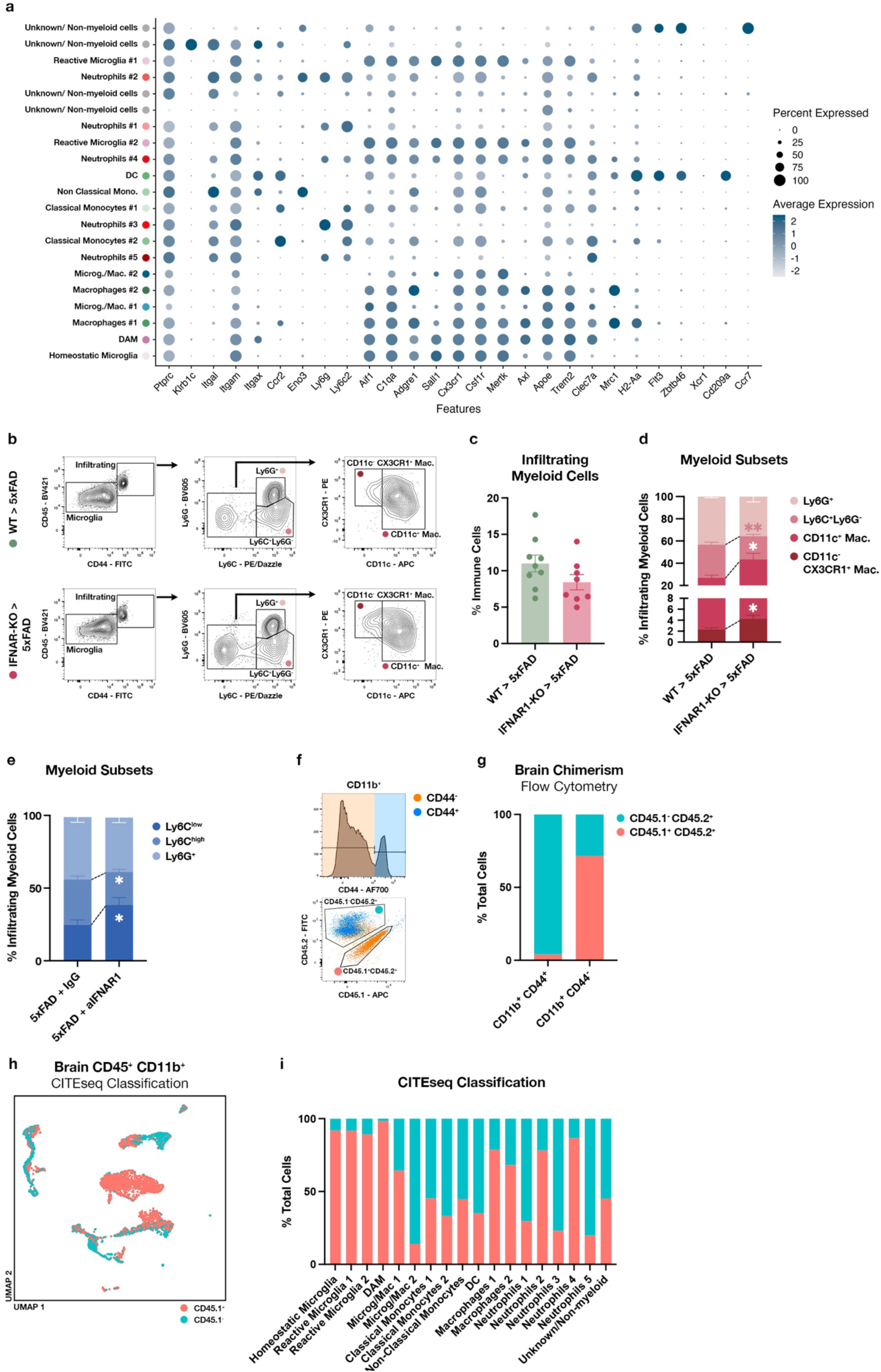
Peripheral IFN-I signaling blockade shifts myeloid cell phenotype in the brain. **a**, Dot plot showing gene markers used for the general annotation of brain myeloid cells analyzed by scRNAseq from 11-month-old WT>5xFAD and IFNAR1-KO>5xFAD chimeric mice (*N* = 4 animals/group; all were females). **b**, Gating strategy and representative contour plots of the flow cytometry analysis of infiltrating myeloid cells in the brains of the 5xFAD chimeric mice. **c**, Infiltrating myeloid cells (CD11b^+^CD45^high^CD44^+^) in the brain of 5xFAD chimeric animals (*N* = 8-9 animals/group). Unpaired t-test. **d**, Proportions of the different subsets of infiltrating myeloid cells in the brains of 5xFAD chimeric mice analyzed by flow cytometry (*N* = 8-9 animals/group). Unpaired t-test for each cell type. **e**, Proportions of the different subsets of infiltrating myeloid cells (CD45^high^CD44^+^CD11b^+^) in the brains of 5xFAD mice treated with IgG or aIFNAR1 monoclonal antibodies (*N* = 10 animals/group). Unpaired t-test for each cell type. **f**, Gating strategy used for the flow cytometry analysis of the peripheral origin of CD11b^+^CD44^+^ cells in the brain of chimeric mice. **g**, Proportions of peripheral (CD45.1^-^CD45.2^+^) or brain resident (CD45.1^+^CD45.2^+^) myeloid cells among the CD11b^+^CD44^+^ and CD11b^+^CD44^-^ cells as determined by flow cytometry (*N* = 3). **h**, UMAP showing the CITEseq classification of CD45.1 positive or negative cells generated by scRNAseq analysis of CD11b^+^B220^-^TCRβ^-^ sorted from the brain of 11-month-old chimeric mice (WT>5xFAD, *N* = 4; IFNAR1-KO>5xFAD, *N* = 4; all were females). **i**, Proportion of CD45.1^+^ (resident-derived) or CD45.1^-^ (BM-derived) cells in the clusters from the scRNAseq analysis of myeloid cells from chimeric mice brains.

**Extended Data Fig. 11:**
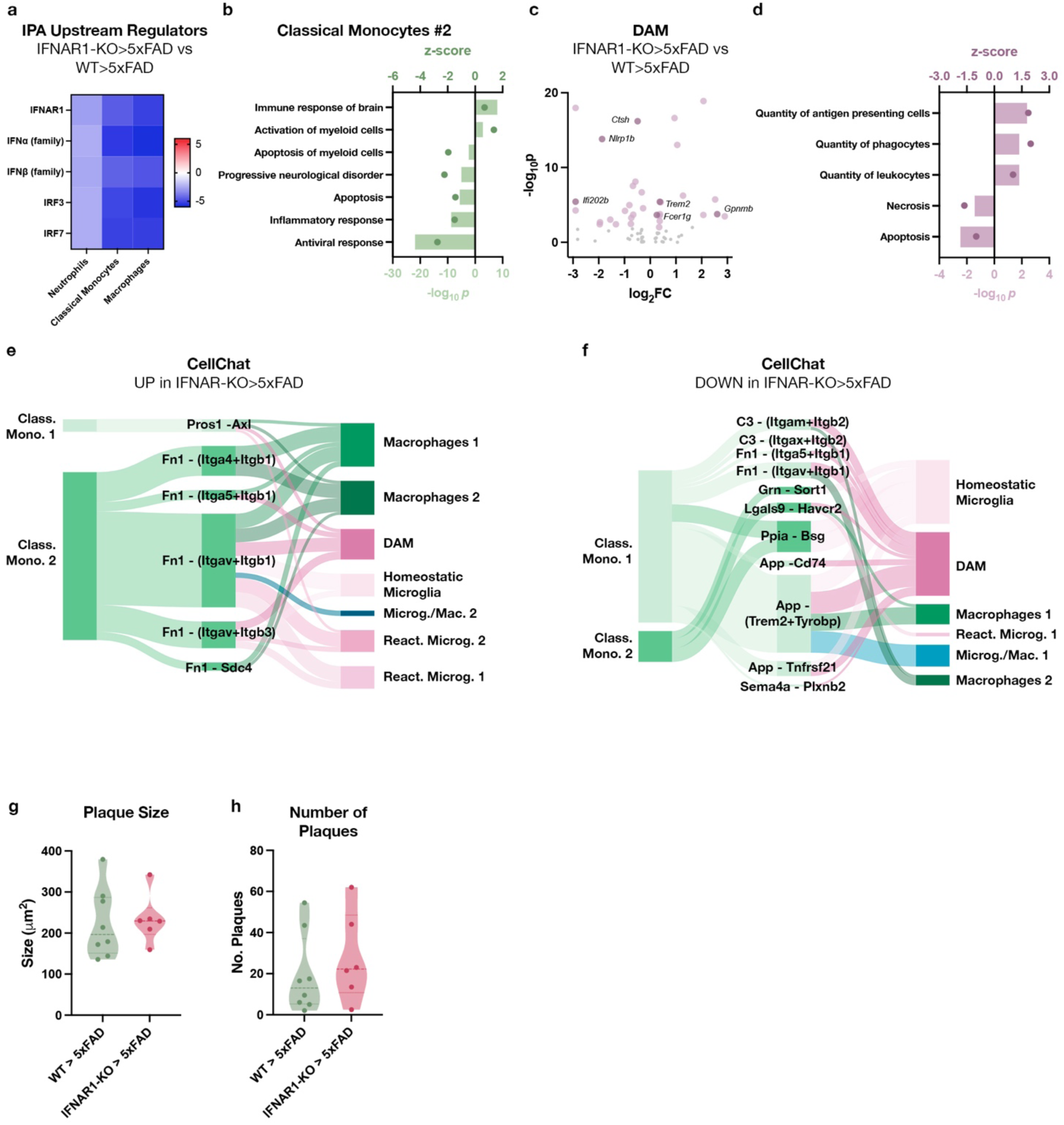
Cell-cell interaction networks are affected in the brains of IFNAR1-KO chimeric 5xFAD mice. **a**, IPA ‘Upstream Regulators’ of infiltrating neutrophils, monocytes and macrophages in the brain of IFNAR1-KO>5xFAD chimeric mice compared to WT>5xFAD chimeric mice. **b**, IPA ‘Diseases and Functions’ predicted from the DEG from classical monocytes cluster #2, selected among the ones with z-score > 1 or < −1, and p-value < 0.01. **c**, DEG in DAM from IFNAR1-KO>5xFAD compared to WT>5xFAD. Genes highlighted in lavender have log_2_FC > 0.25 or < −0.25 and p-value < 0.01. **d**, IPA ‘Diseases and Functions’ predicted from the DEG from DAM, selected among the ones with z-score > 1 or < −1, and p-value < 0.01. **e**, Sankey plot of cell-cell interactions increasing in the brains of IFNAR1-KO>5xFAD mice compared to WT>5xFAD mice. **f**, Sankey plot of cell-cell interactions decreasing in the brains of IFNAR1-KO>5xFAD mice compared to WT>5xFAD mice. **g**, Histological quantification of average amyloid plaque size in the CA1 of 5xFAD chimeric mice (*N* = 6-8 animals/group). **h**, Histological quantification of the number of amyloid plaques in the CA1 of 5xFAD chimeric mice (*N* = 6-8 animals/group).

